# Brain biomechanics governs mitotic fidelity of embryonic neural progenitors

**DOI:** 10.1101/2025.07.31.667957

**Authors:** V. Marthiens, C. Basto, L. Jawish, M. Malosse, D. Krndija, A.S. Mace, E. Laigna, C. Peyret, T. Pan, O. Zajac, L. Perrin, E. Logarinho, C. Villard, R. Basto

**Author notes:** These authors contributed equally to this work (C. Basto, L. Jawish, M. Malosse).

## Abstract

Accurate chromosome segregation is essential to maintain genetic stability and prevent the onset of diseases such as developmental disorders, infertility, and cancer. While many intrinsic factors and processes involved in mitosis have been extensively characterized, less is known about how extrinsic factors and tissue properties contribute to mitotic fidelity. In this study, using both *in vivo* and *ex vivo* systems, in combination with pharmacological perturbations and high-resolution microscopy, we investigated mitosis in apical radial glial (aRG) cells—the primary neural progenitors in the developing mammalian brain. We found that the high cell density typical of early neurogenic stages enhances microtubule polymerization rates from the spindle poles, thereby influencing chromosome segregation. Mechanistically, our results indicate that cortical actin and elevated cortical tension function as sensors of biomechanical stress during mitosis. These findings identify biomechanics as a threat to mitotic fidelity in the embryonic brain.

## Introduction

The mechanisms controlling chromosome behaviour during mitosis have long been studied outside of native tissues in mammals. However, whether tissue properties may influence mitotic parameters including chromosome segregation fidelity has remained unknown. The embryonic mammalian brain is particularly attractive to address this question. First, this tissue is prone to errors in chromosome segregation errors in adverse circumstances (Lizarraga *et al*, 2010; McIntyre *et al*, 2012; Marthiens *et al*, 2013; Marjanović *et al*, 2016; Shi *et al*, 2019; Viais *et al*, 2021), even if the nature and frequency of resident aneuploid cells remains a matter of debate (Rehen *et al*, 2005; McConnell *et al*, 2013; Cai *et al*, 2014; Knouse *et al*, 2014). Second, mutations in mitotic genes have been described in human neurodevelopmental disorders, like human MicroCephaly Primary Hereditary (MCPH), which predominantly disrupt brain size while sparing other tissues of the organism (Thornton & Woods, 2009; Kaindl *et al*, 2010; Jayaraman *et al*, 2018; Naveed *et al*, 2018; Farcy *et al*, 2023). Third, it has been previously reported that apical radial glial cells (aRG), the population of neural progenitors at the origin of neurons and glial cells, are more susceptible to undergo chromosome mis-segregation at early developmental stages. (Marthiens *et al*, 2013; Vargas-Hurtado *et al*, 2019; Marthiens & Basto, 2020). This suggests that embryonic brain tissue properties may evolve during neurogenesis and favour the generation of aneuploidy at certain developmental stages. Importantly, aneuploidy in the developing brain results in premature neuronal differentiation and/or p53-dependent apoptosis, linking chromosome mis-segregation with brain size reduction through depletion of the progenitor pool (Lizarraga *et al*, 2010; Marthiens *et al*, 2013; Gogendeau *et al*, 2015; Marjanović *et al*, 2016; Pilaz *et al*, 2016; Shi *et al*, 2019).

During development, the cerebral cortex is submitted to intense tissue remodelling during a very short time window, as neuron production must be achieved at birth (Paridaen & Huttner, 2014; Casas Gimeno & Paridaen, 2022; Villalba *et al*, 2021). The biogenesis of the mammalian neocortex takes place under a high load of biomechanical constraints, imposed by progenitor cell density, interkinetic nuclear migration (where nuclei shuttle up and down within the neuroepithelium during the cell cycle) and neuron migration towards the cortical layers, to cite a few. Whether neural progenitor mitosis is sensitive to such changes imposed by the embryonic brain environment has not been investigated so far. Here, we have characterized the biomechanical properties of the mouse developing brain in which aRG cells that give rise to all populations of neurons and glial cells divide during neurogenesis. We found that at early stages, aRG cells undergo mitosis in a confined environment, favouring the assembly of mitotic spindles that fail to segregate chromosomes correctly. Mechanistically, we show that aRG cells submitted to the compressive stress imposed by their neighbors contain an actin rich cortex that increase the frequency of chromosome mis-segregation by promoting microtubule (MT) polymerization from the spindle poles. These results link cell density and cortical membrane tension with mitotic fidelity in the mammalian brain.

## Results

### Progenitor shape, volume and membrane properties change along with cell density in the developing mouse neuroepithelium

During mouse brain development, embryonic neural stem cells, also called apical radial glia (aRG) due to their elongated shape and glial features, give rise to all the neuronal and glial populations that compose the brain (Villalba *et al*, 2021). aRG keep bipolar contacts *via* thin processes - apical and basal processes - with the ventricle and the *basal lamina,* respectively (Figure 1A) (Taverna & Huttner, 2010; Arai & Taverna, 2017). aRG cells undergo mitosis close to the ventricle and we have shown that mitotic spindle morphology changes during brain development (Vargas-Hurtado *et al*, 2019). To probe whether morphometrics of mitotic aRG cells also change during development, we analysed the ventricular zone from *en face* view at the peak of aRG expansion at E13.5 in comparison with E16.5.

**Figure 1:**
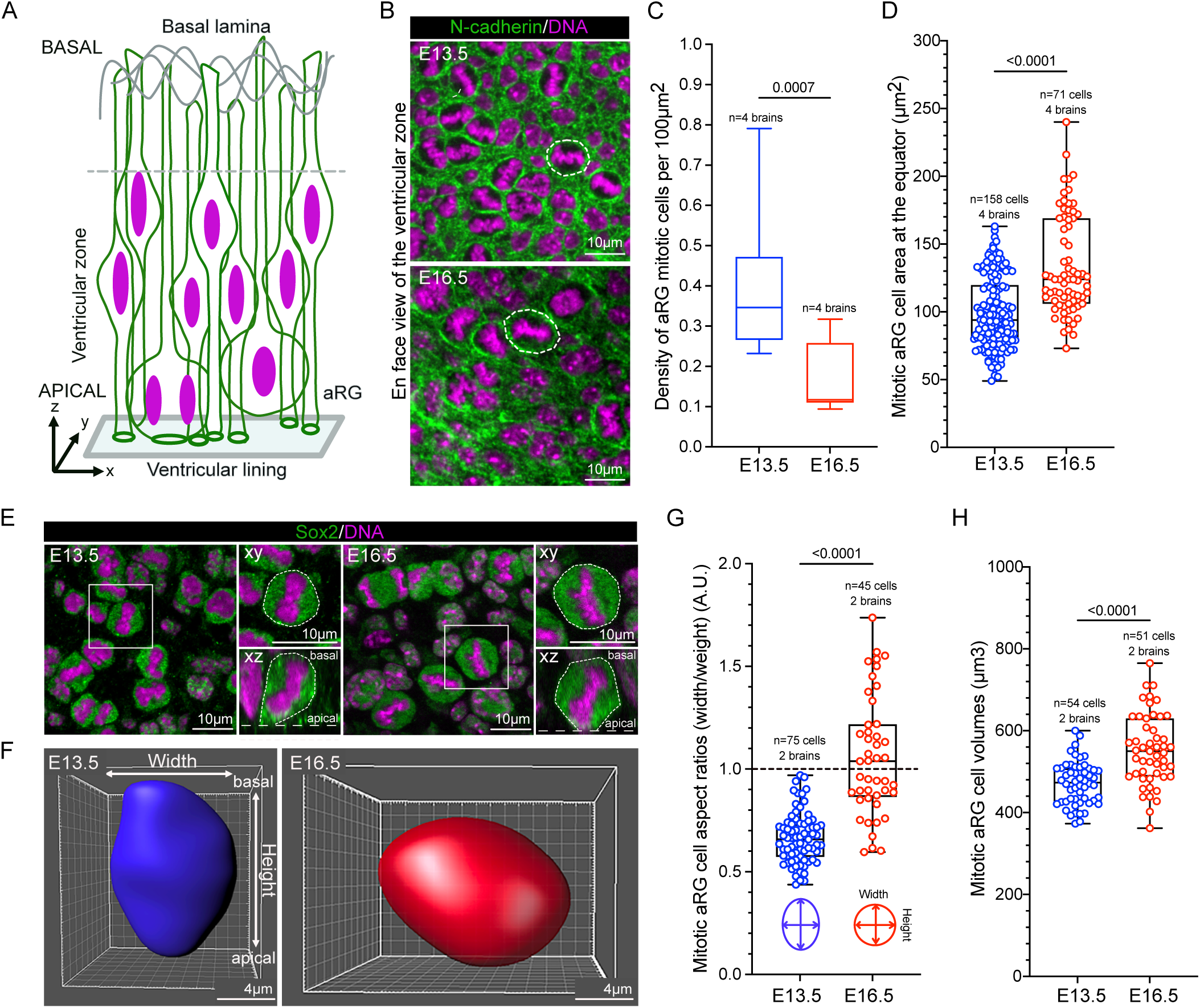
Changes in aRG cell mitotic morphometrics during neurogenesis. (A) Schematic diagram depicting apical Radial Glia (aRG) apico-basal organization in the neuroepithelium. The nucleus in magenta and the plasma membrane in green. (B) Representative pictures of *en face* views of whole-mount cerebral cortex explants at E13.5 (upper panel) and E16.5 (bottom panel) showing the nucleus (magenta) and plasma membrane labelled with N-cadherin antibodies (green). White dashed lines highlight the contour of mitotic cells for comparison. (C-D) Box and whiskers dot plots of mitotic aRG cell density (C) and aRG cell equatorial area (D) at E13.5 and E16.5. (E) Representative pictures of *en face* views (xy) and lateral (xz, with apical side indicated by the dashed white line at the bottom) views of whole-mount cerebral cortex explants labelled with Sox2 antibodies (green). DNA in magenta. (F) 3D-reconstruction of E13.5 (blue shape, left panel) and E16.5 (red shape, right panel) aRG cell volumes in whole-mount cerebral cortex explants. (G) Box and whisker dot plot of aRG mitotic cell aspect ratios (G) as indicated. Note that the height is measured along the apico-basal axis and the width perpendicular to it. The black dotty line point to an aspect ratio of 1, which corresponds to a round shape mitosis. (H) Box and whisker dot plot of aRG mitotic cell volumes in mitosis as indicated. Data were submitted to normality and lognormality tests for the choice of appropriate statistical test. Statistical significance was assessed with Mann-Whitney tests (C-D and H) or an unpaired t-test (G).

aRG cell density decreased during neurogenesis as illustrated by a three-fold decline in apical process density at the ventricular lining (Figures S1A-B) and apical density of mitotic cells (Figures 1B-C). Such a decline in apical cell density was accompanied by an increase in the area of both apical processes (Figure S1C) and mitotic cells at the equator (Figure 1D), which are good indicators of a decrease in spatial constraints in the ventricular zone as neurogenesis progresses. Moreover, lateral constraints imposed by neighboring cells in the ventricular zone appeared to impact both mitotic cell shape, as evidenced by a change in aRG aspect ratios (ratios of cell width over height) from an elongated shape at E13.5 (mean aspect ratio at 0.65) to a circular shape at E16.5 (mean aspect ratio at 1) (Figures 1E-G) and a concomitant increase in cell volume (Figures 1H and S1D).

Altogether, our results show a decrease in mitotic aRG density at the ventricular zone accompanied by cell rounding and volume increase during neurogenesis.

### Mechanical confinement by neighboring cells promotes mitotic errors in aRG

The changes observed *in vivo* in the mouse brain between E13.5 and E16.5 correlate with differences in mitotic fidelity (Vargas-Hurtado *et al*, 2019). To evaluate whether there is a causal relationship between aRG density and chromosome segregation errors, we established an *ex vivo* approach to experimentally manipulate neighboring cell density of aRG mitotic cells. We developed a protocol where aRG cells were dissociated from E13.5 mouse dorsal telencephalon to establish primary cell cultures (Figures S2A-B). We expanded these cultures either as 3D floating spheres - referred to here as 3D aRG or plated them on fibronectin substrates - called 2D aRG (Figure 2A). Upon mitotic entry, 2D aRG like other cells in culture, de-adhere before rounding up (Taubenberger *et al*, 2020) and undergo mitosis without lateral cell-cell contacts (Figure 2B). The morphometrics of aRG mitosis differed in the two experimental set-ups, with 2D aRG mitotic cells exhibiting a more circular shape (as evidenced by the measure of aspect ratios, i.e. cell height over cell length) (Figure 2C) and increased cell volume (Figure 2D), as compared to 3D aRG mitotic cells. The presence of neighboring cells around mitosis *in vitro* is therefore sufficient to influence cell shape and volume, in agreement with our *in vivo* observations in the mouse developing brain.

**Figure 2:**
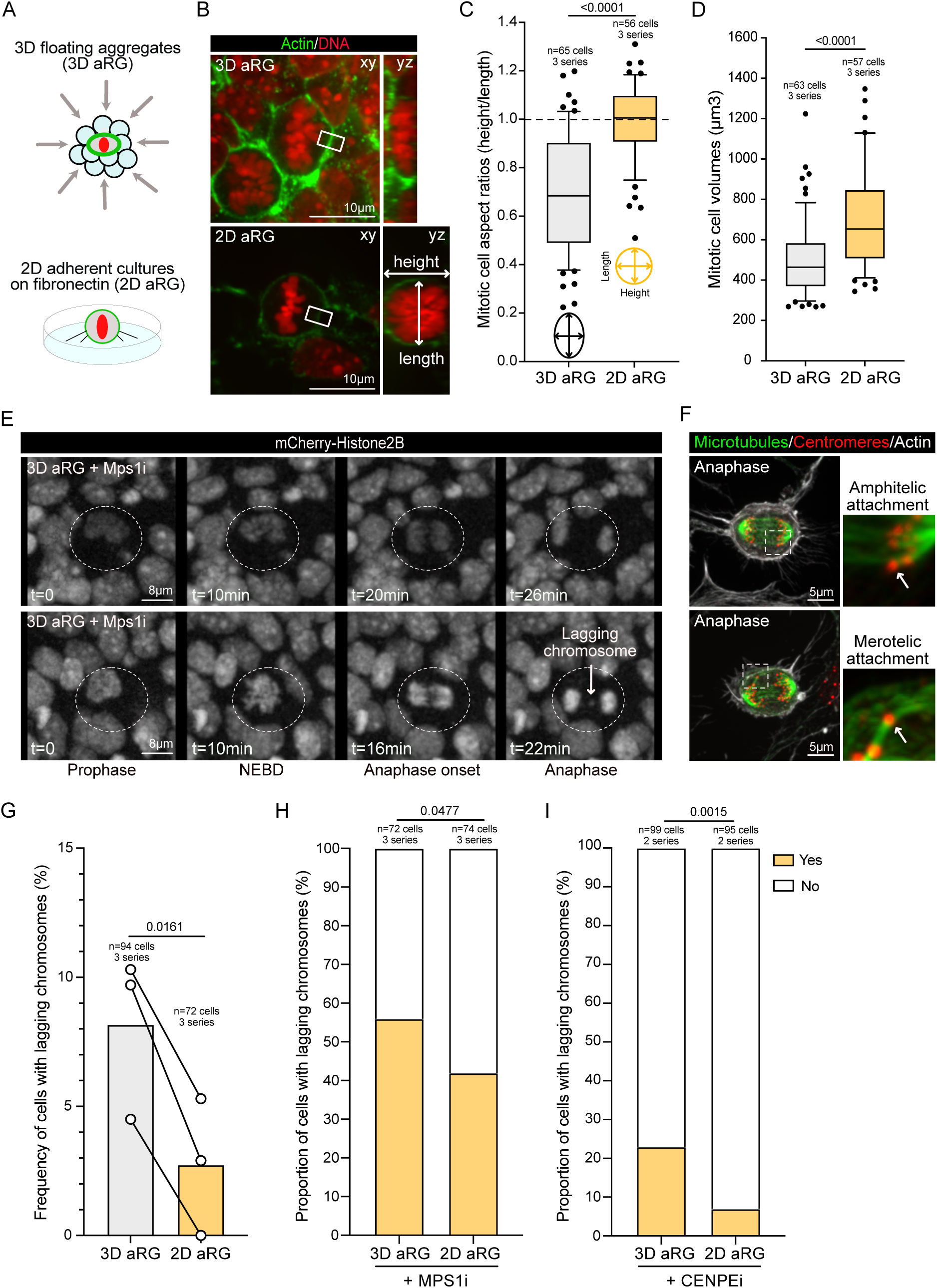
The frequency of chromosome segregation errors is influenced by the presence of neighboring cells. (A) Schematic diagram showing 3D (top) and 2D (bottom) experimental set-ups for aRG primary cell cultures. (B) Representative pictures with top (left panel, xy) and side (right panel, yz) views of 3D (top) or 2D (bottom) aRG mitosis showing actin (green) and DNA (red). (C-D) Dot plot bar graphs showing mitotic aRG cell aspect ratios (C) and volumes (D) as indicated. (E) Stills of time lapse movies of aRG cells expressing mCherry-histone2B to visualize DNA (in grey) treated with MPS1i from prophase to anaphase. The last panel on the right shows anaphases with (bottom) or without (top) lagging chromosomes (white arrow). (F) Representative pictures of aRG cells in anaphase labelled with antibodies that recognize MTs (green), centromeres (red) and actin (grey). The dashed squares show the higher magnification insets on the right to show different types of kinetochore-MT attachments as indicated. (G) Graph bar showing the percentage of aRG cells with lagging chromosomes scored by live imaging as indicated. (H-I) Graph bars showing the proportion of cells with lagging chromosomes as indicated, in the presence of MPS1i (H) or CENPEi (I). Data were submitted to normality and lognormality tests for the choice of appropriate statistical test. Statistical significance was assessed with non-parametric unpaired Mann-Whitney tests (C-D), parametric paired t-test (G) and chi-square on normalized contingency data (H and I).

Based on these observations, we next compared mitotic fidelity in aRG cells with and without cell neighbors −3D and 2D conditions, respectively. To evaluate the frequency of chromosome segregation errors, we used a mCherry-Histone2B (mCherry-H2B) expressing mouse line to score, by live imaging, the frequency of lagging chromosomes in anaphase (Figure 2E) resulting from merotelic attachments (Figure 2F). The frequency of chromosome mis-segregation was more than two times higher in 3D than in 2D aRG cultures, as illustrated by either comparing the frequency of lagging chromosomes *per* set of experiment (Figure 2G) or by displaying the proportion of cells exhibiting lagging chromosomes in three sets of experiments (Figure S2D). Importantly, these differences cannot be explained by differences in mitotic timing (Figure S2C).

In order to test whether it was possible to emphasise the increased frequency of lagging chromosomes in 3D aRG mitotic cells as compared to 2D cells, we used the spindle assembly checkpoint (SAC) MPS1 inhibitor (MPS1i) (Santaguida *et al*, 2010; Tardif *et al*, 2011). We used low concentrations of MPS1i (25nM) in an acute manner (maximum of 2.5h-to-3h time lapse recording in the presence of the drug). This concentration was sufficient to inactivate the SAC, as illustrated by a noticeable decrease in the time spent by aRG cells in mitosis (from 21min in control to 10-12min in MPSi) (Figure S2E). Moreover, as expected, the percentage of anaphases with lagging chromosomes was increased as compared to DMSO (Figure S2F). Strikingly, and in agreement with our previous observations under unchallenged conditions, the frequency of chromosome mis-segregation was still higher in 3D *versus* 2D aRG cells treated with MPS1i (Figures 2H and S2F-G). It is worth noticing that MPS1i treatment caused a more severe reduction of mitotic timing in aRG cells in 2D than in 3D (Figure S2E) (mean of 10min *versus* 12min), likely reflecting differential access to the drug between the two experimental set-ups. Further, we used the Centromere Protein E (CENP-E) kinesin inhibitor (CENPEi). CENP-E is a component of the kinetochore *fibrous corona* that promotes chromosome congression and kinetochore-MT stability (Weaver *et al*, 2003; Gudimchuk *et al*, 2013; Vitre *et al*, 2014). Similar to MPS1i treatment, CENPEi resulted in a higher frequency of chromosome segregation errors in aRG cells cultured in 3D than in 2D (Figures 2I and S2H-I).

Collectively, our data indicate that the presence of neighboring cells can influence the rate of chromosome mis-segregation in both unchallenged and challenged conditions.

## Spatial constraints and cell adhesion mimicking cell-cell contacts increase astral microtubule density

We have previously reported that aRG spindles are particularly enriched in astral MTs when they are prone to mitotic errors in E13.5 cerebral cortex (Vargas-Hurtado *et al*, 2019). We therefore hypothesized that the biomechanical constraints faced by mitotic cells in the presence of cell neighbors (3D aRG cultures) may favour the nucleation or stabilization of astral MTs. To test this possibility, and in order to mimic the lateral constraints imposed to aRG mitotic cells in the brain, we seeded primary cultures of E13.5 aRG dissociated from the dorsal telencephalon (Figures S2A and S2B) on top of a polydimethylsiloxane (PDMS)-based device containing trenches of 10µm in height and between 8 to 12µm in width (Figure 3A). This experimental set-up imposed passive lateral spatial constraints to isolated aRG cells as they undergo mitotic rounding. To facilitate cell seeding and survival, the trenches were coated with fibronectin at the bottom by using the technique of in-mold patterning (Yamada et al, 2016). Furthermore, the wells were coated with N-cadherin-Fc chimera (NcadFc) to mimic cell-cell adhesions. The levels of constraints imposed were evaluated by aRG cell aspect ratios as a measure of cell shape changes (Figures 3B and 3C). This set up allowed us to use cells seeded outside of the wells as an internal control of cells without any constraint (Figure 3A). A distinction between low (trench width above 9.4µm) and high (trench width below 9.4µm) levels of constraint was considered (Figures 3B and 3C). Increasing constraint levels was sufficient to increase the number and length of astral MTs in a stepwise manner, which was further exacerbated by mimicking cell-cell adhesion (Figure 3D and E). It is nonetheless interesting to note that the length of astral MTs reached a plateau in conditions of high constraint, likely because of cell size limit.

**Figure 3:**
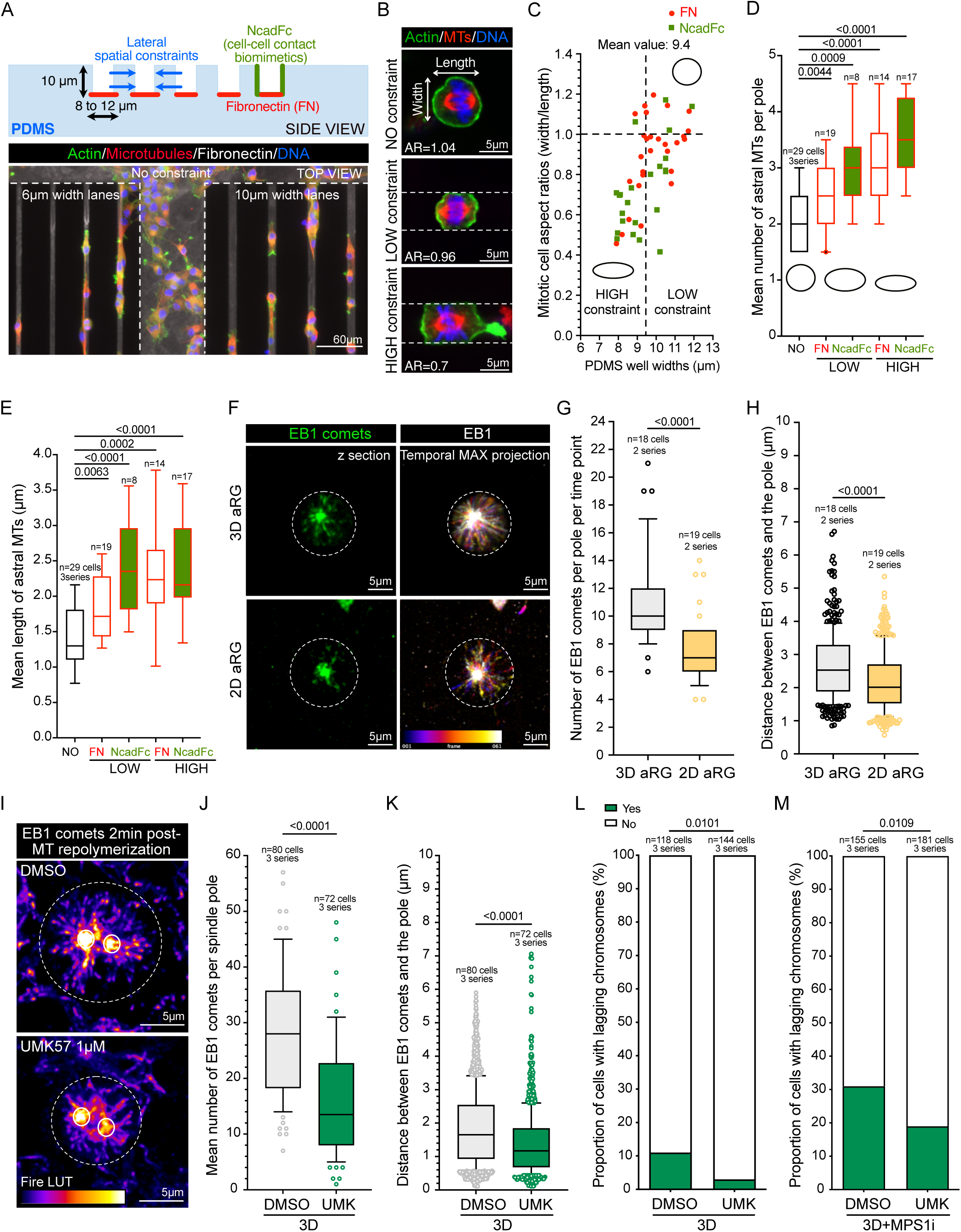
The presence of neighboring cells modulates the rates of spindle pole MT polymerization. (A) Top, schematic diagram of side view of the PDMS device used to spatially constrain mitotic aRG cells. Cells were plated on fibronectin (red) in trenches of different widths ranging from 6 to 12μm. Certain trenches were coated with the N-cadherin biomimetic Ncad-Fc (green). Bottom, representative picture of E13.5 aRG cultures constrained in fibronectin-coated lanes (grey) and visualized with actin (green), MTs (red) and nuclei (blue). Cells were either plated outside of the trenches (no constraint), in trenches of 6μm (left side) or 10μm (right side) widths. (B) Higher magnifications of aRG cells in metaphase outside of constraint (NO constraint-aspect ratio (AR) 1.04), under low (AR: 0.96) or high (AR: 0.7) constraints. (C) Scatter dot plot depicting aspect ratios of individual aRG cells in metaphase according to PDMS width. (D-E) Box and whiskers plot representing the mean number (D) and length (E) of astral MTs in the indicated conditions. Data are provided for cells outside of constraints (NO, empty black boxes), for cells plated on FN in the PDMS wells (empty red boxes), for cells in wells coated with Ncad-Fc (red boxes filled in green). (F) Representative pictures of aRG cells in prometaphase in 3D (top) or 2D (bottom) cultures showing EB1-positive MT (+) tips polymerized from one pole represented at one time point (left panels, EB1 in green) or as a temporal maximum projection during a 1min long MT repolymerization movie with acquisitions taken every second (right panels). False colours represent the time when the EB1 comets were first detected during the 1min recording. (G-H) Box and whiskers plots showing the number of EB1 comets polymerized per pole at a given time point (G) and the maximal distance travelled by individual EB1 comets from the spindle pole (H) as indicated. (I) Representative images of aRG cells in prometaphase at t=2min after the start of MT repolymerization showing the accumulation of EB1-positive MT (+) tips (false colours, with the fire LUT representing the levels of EB1 fluorescence intensity) around the two spindle poles as indicated. Note the asymmetry in MT polymerization rate between the two spindle poles. (J-K) Box and whiskers plots showing the cumulative number of EB1 comets polymerized per pole (J) and the distance travelled by individual EB1 comets from the spindle poles (K) during 2min of MT repolymerization as indicated. (L-M) Graph bars showing the percentage of cells displaying (green boxes) or not (white boxes) lagging chromosomes in time-lapse experiments as indicated. Data were submitted to normality and lognormality tests for the choice of appropriate statistical test. Statistical significance was assessed with Mann-Whitney tests (D, G-H, J-K), unpaired t test (E) and Fisher’s exact test (L-M).

We concluded that the presence of lateral constraints is sufficient to increase the frequency and length of astral microtubules in mitotic spindles.

### Spatial constraints (but not cell-cell adhesion *per se*) enhance spindle pole MT polymerization

The increase in astral MT number and length under constraint suggested promotion of MT polymerization capacity from the spindle poles. We investigated MT polymerization activity by tracking plus-end EB1-positive comets as a readout of actively polymerizing MT plus tips in mitosis (Muroyama & Lechler, 2017; McHugh & Welburn, 2023). Since MT polymerization activity is asymmetric between the two spindle poles, we plotted the data of the pole displaying the highest activity. Comparison between 3D and 2D conditions revealed higher MT polymerization rates in 3D aRG cells, illustrated by 1.5-fold higher number of EB1 comets polymerized per time point (Figures 3F and 3G). Moreover, the distance travelled by EB1 comets from the spindle pole, which reflects MT polymerization persistence (Muroyama & Lechler, 2017; Roostalu *et al*, 2020), was also higher (Figure 3H). Our observations were similar after MT depolymerization-repolymerization assays performed on a short time scale (1.5min repolymerization, Figure S3A), evaluating the same parameters for individual EB1 comets (Figures S3B-C).

We next determined whether the density of cell-cell adhesion was triggering the changes in spindle pole MT polymerization capacity. 2D aRG cultures were plated on the cell-cell contact biomimetic Ncad-Fc chimera instead of fibronectin (Figures S3D-F). Interestingly, while mitotic rounding was affected by the presence of Ncad-Fc as a substrate, as illustrated by the flattened aspect of the cells at the interface with the substrate (Aspect Ratio below 1, as compared to the FN control substrate) (Figure S3D), the number and length of actively polymerizing MTs were not noticeably affected (Figures S3E-F). These observations support the conclusion that N-cadherin-dependent cell-cell contacts are not promoting spindle pole MT polymerization activity but are more likely stabilizing the population of astral MTs at the plasma membrane (Figures 3D and 3E).

Altogether, our results show that during mitosis in aRG cells, biomechanical constraints imposed by neighboring cells can modulate the rate of MT polymerization at the spindle poles, with a poor contribution of cell-cell adhesion *per se*.

### Biomechanical constraints induce spindle pole polymerization activity promoting chromosome mis-segregation

MT dynamic instability is achieved by a balance between MT polymerization and depolymerization events (Mitchison & Kirschner, 1984). Depolymerases such as MCAK/Kif2C play key roles in mitotic spindle assembly and chromosome segregation in many cell types (Stout *et al*, 2011; Tanenbaum *et al*, 2011; Walczak *et al*, 2016; van Heesbeen *et al*, 2017; McHugh & Welburn, 2023). We functionally tested whether counteracting the high polymerization activity typical of 3D aRG mitosis could lower the frequency of chromosome mis-segregation. To this end, we used the small-molecule agonist of MCAK motor activity - UMK57 - which decreases MT polymerization and consequently the frequency of lagging chromosomes in aged human fibroblasts (Barroso-Vilares *et al*, 2020). In 3D aRG cells, this treatment had the expected effect of decreasing the number and length of actively polymerizing MTs (Figures 3I-3K). Importantly, a decrease in the frequency of lagging chromosomes (Figures 3L and S3H), without differences in mitotic timing (Figure S3G) was observed. Moreover, UMK57 treatment was also sufficient to reduce the rate of lagging chromosomes in 3D aRG cells challenged with MPS1i (Figures 3M and S3J), while the combination of MPS1i and UMK57 appears to slightly delay mitotic progression (Figure S3I).

We concluded that in mitotic aRG cells, biomechanics can increase chromosome mis-segregation by influencing spindle pole MT polymerization rates.

### Mechanical stress enhances MT polymerization and promotes chromosome mis-segregation

Our findings support a causal relationship between biomechanical constraints, MT polymerization rates and mitotic fidelity in aRG cells (Figures 3D-M, Figure S3). We next investigated the outcome of imposing constraints on 2D aRG cultures, to overcome the lack of cell-cell contacts during mitosis. We used a biocompatible osmolyte of high molecular weight - dextran-that is diluted in the culture medium (Monnier *et al*, 2016; Dolega *et al*, 2021). Dextran exerted a mechanical pressure by counteracting the force required for mitotic cells to round-up, as illustrated by their elongated shape at the interface with the substrate (Figures 4A-B). Importantly, dextran did not alter their circular shape at the equator and therefore preserved the space required to accommodate the bipolar spindle and the metaphase plate (Figure 4A) (Lancaster *et al*, 2013). Strikingly however, this was sufficient to increase spindle pole MT polymerization (Figures 4C-E) and the frequency of lagging chromosomes in the absence (Figures 4F and S4B) or presence of MPS1i (Figures 4G and S4D) without affecting mitotic timing (Figures S4A and S4C).

**Figure 4:**
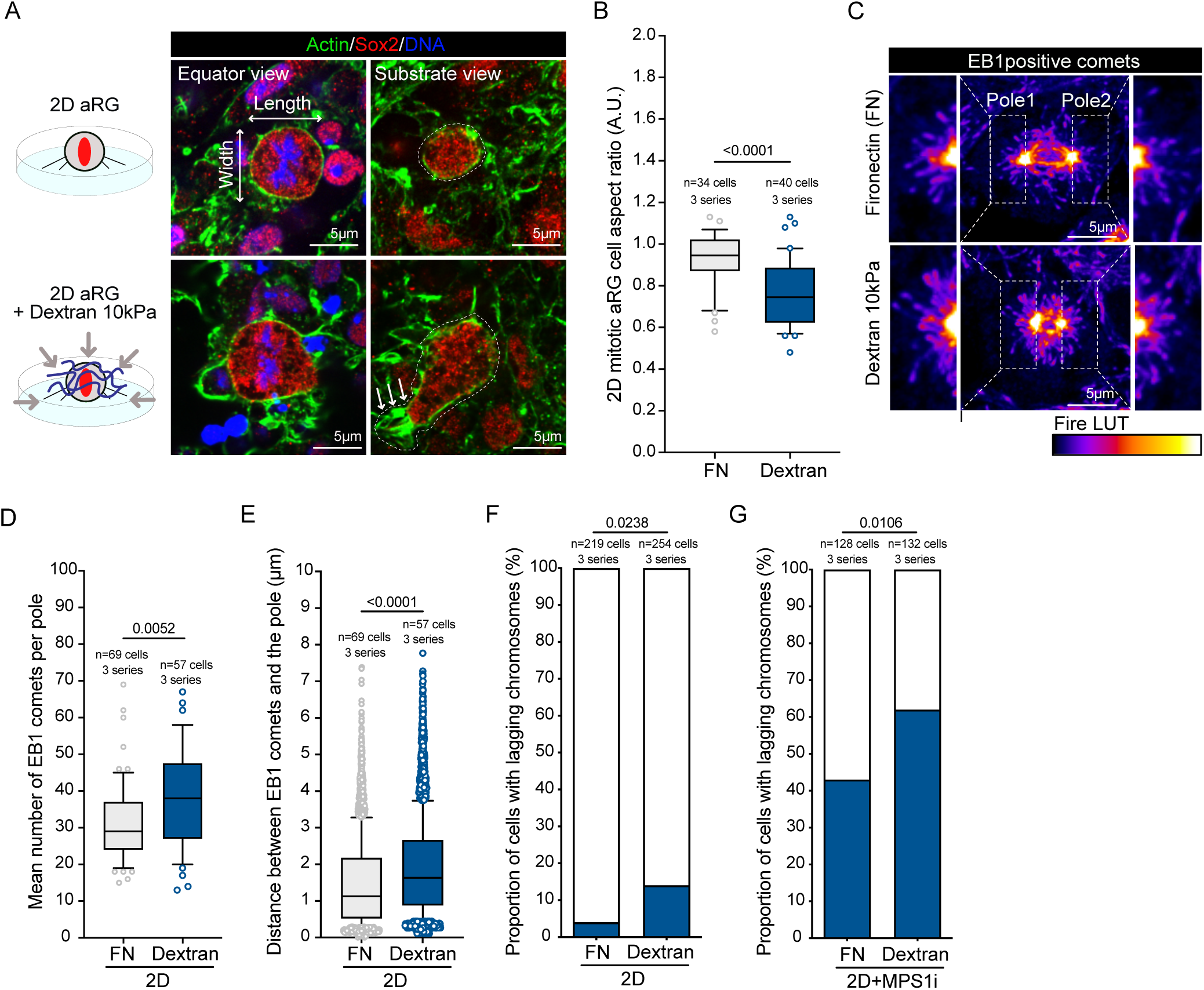
Compression of 2D aRG cells is sufficient to increase MT polymerization and chromosome mis-segregation rates. (A) On the left, schematic diagram of 2D aRG cells with (bottom panels) or without (top panels) dextran at a concentration which imposes a compression of 10kPa. On the right, representative images of equatorial views (left panels) or substrate view (right panels) of aRG cells in metaphase showing actin (green), Sox 2 (red) and DNA (blue). Small white arrows point to retraction fibres. (B) Box and whiskers plot showing 2D aRG aspect ratios evaluated at the level of the substrate as indicated. (C) Representative pictures of EB1-positive comets from aRG cells in metaphase. High magnifications of the two spindle poles are provided as left and right insets. The fire LUT was used to represent the levels of EB1 fluorescence intensity. (D-E) Box and whiskers plot showing the mean number per pole (D) and distance travelled from the pole of EB1 comets (E) in the absence (Fibronectin substrate only, FN) or presence of dextran as indicated. (F-G) Graph bars showing the frequency of lagging chromosomes as indicated. Data were submitted to normality and lognormality tests for the choice of appropriate statistical test. Statistical significance was assessed by Mann-Whitney tests (B, D-E) and Fisher exact test on normalized contingency data (F and G).

Our results show that the compressive stress imposed by 3D constraints when cells undergo mitosis in multicellular environments -such as the aRG spheroids-dictates spindle pole MT polymerization capacity and consequently chromosome segregation accuracy.

### Cortical actin senses biomechanical compression in mitotic aRG cells

The mechanism for sensing compression levels as an indicator of neighboring cell density in 3D aRG in mitosis remains unknown. To identify possible cell sensors, we compared the levels of cortical actin, which contributes to mitotic cortical tension (Stewart *et al*, 2011; Chugh *et al*, 2017; Taubenberger *et al*, 2020) in E13.5 and E16.5 brains. We also characterized phosphorylated Ezrin Radixin and Moesin (pERM) proteins at the cell cortex as active pERM proteins link the plasma membrane and actin filaments to provide plasma membrane rigidity (Carreno *et al*, 2008; Kunda *et al*, 2008; Machicoane *et al*, 2014; Taubenberger *et al*, 2020). Interestingly, both cortical actin (Figures 5A-C) and p-ERM levels (Figures S5A-C) were higher at E13.5 as compared to E16.5. To determine whether the observed differences in actin and pERM recruitment reflected changes in cortical membrane tension, we measured the initial velocity recoil speed after laser-cuts of aRG cells to obtain relative values of cortical tension (Farhadifar *et al*, 2007; Rujano *et al*, 2013; Krndija *et al*, 2019). We used embryonic brains endogenously expressing LifeAct-GFP (Figure 5D). We observed a higher velocity of membrane recoil, indicative of higher cortical membrane tension in E13.5 brains as compared to E16.5 brains (Figures 5E-F).

**Figure 5:**
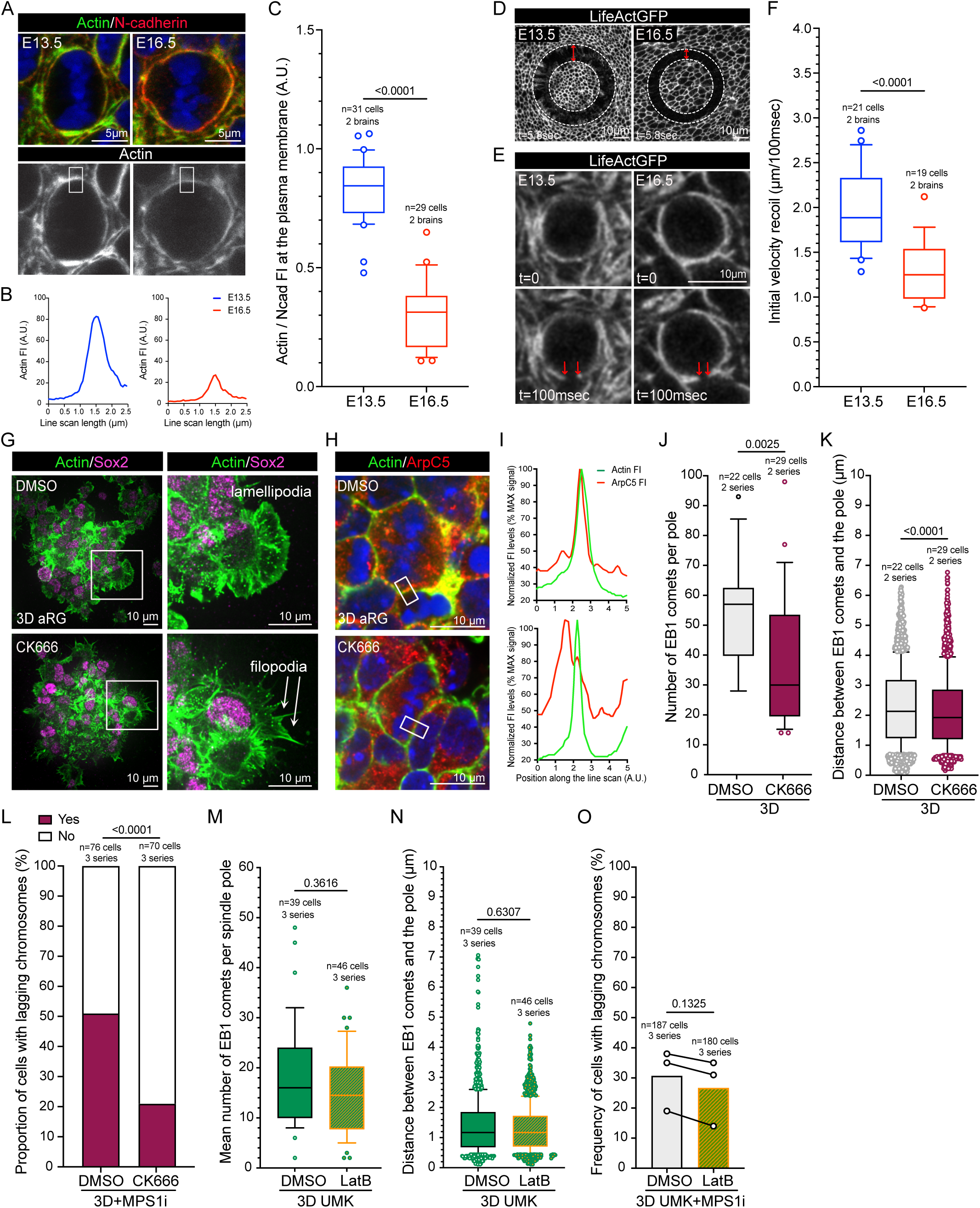
Cortical actin modulates MT polymerization and chromosome mis-segregation rates. (A) Representative images of aRG cells in mitosis from embryonic cerebral cortices at E13.5 (left) and E16.5 (right) showing actin (green), N-cadherin (red) and DNA (blue). Actin is shown in the bottom panels in grey. The white rectangles in (A) point to the regions chosen for line scan analysis of actin fluorescence intensity provided in (B). (C) Box and whiskers plot showing actin over N-cadherin fluorescence intensity (FI) measures as indicated. (D) Representative *en face view* images of the ventricular lining showing LifeAct-GFP boundaries (in grey) after a circular shape laser cut. The red arrows measure the distance after 5.8 seconds of recoil velocity. (E) Representative images of mitotic aRG cells from embryonic cerebral cortices expressing LifeAct-GFP at the plasma membrane before (t=0, upper panels) and 100 milliseconds (msec, lower panels) after laser cut. The two red arrows indicate the distance of recoil. (F) Box and whisker plot showing initial velocity recoil after laser cut of mitotic aRG plasma membrane. (G) On the left, representative images of 3D aRG cultures labelled to show actin (green) and the stem cell marker Sox2 (magenta). On the right, high magnifications of the white squared regions highlighted on the left panels depicting lamellipodia after DMSO (top panels) and filopodia after CK666 (bottom panels) treatments. (H) Representative pictures of 3D aRG cultures showing ArpC5 localization (red) and actin (green) as indicated. (I) Actin (green line) and Arpc5 (red line) FI line scans of the plasma membrane. Positions of the line scans are depicted as white rectangles in (H). (J-K) Box and whiskers plots showing the number of EB1 comets polymerized per pole (J) and the distance of individual EB1 comets travelled from the spindle pole (K) after 2min of MT repolymerization as indicated. (L) Graph bar showing the proportion of cells with lagging chromosomes as indicated. (M-N) Box and whiskers plots showing the number of EB1 comets polymerized per pole (M) and the distance of individual EB1 comets from the spindle pole (N) after 2min of MT repolymerization following 1H DMSO (grey boxes) or LatB (orange-green boxes) pre-treatment in the presence of the MCAK agonist UMK57 (UMK). (O) Graph bar showing the frequency of lagging chromosomes scored in 3D aRG cells as indicated. Data were submitted to normality and lognormality tests for the choice of appropriate statistical test. Statistical significance was assessed with Mann-Whitney tests (F, J, K, M and N), an unpaired t test (C), a paired t test (O) and a Chi square test (L).

We hypothesized that cortical actin may be a key parameter to sense compression levels as cortical membrane tension is driven by the length of actin filaments in mitosis (Chugh *et al*, 2017). To test this possibility, we first used latrunculin B (LatB) at very low doses (50nM) in an acute manner (1h to 2.5h of incubation). LatB binds actin monomers and therefore prevents actin filament polymerization (Spector *et al*, 1983). In 3D aRG configurations, LatB treatment resulted in a decrease in cortical actin recruitment (Figure S5D) and decreased cortical membrane tension after laser cut experiments (Figures S5E-F). In agreement with the literature, mitotic cell volumes remained unaltered after LatB treatment (Figure S5G) (Zlotek-Zlotkiewicz *et al*, 2015). Importantly, MT depolymerization-repolymerization assays and analysis of EB1 comet behaviour revealed a decrease in both the number and length of polymerizing MTs from the spindle poles (Figures S5H-I). Furthermore, in addition to decreasing polar MT polymerization, a decrease in the frequency of lagging chromosomes in MPS1i (Figure S5K) or CENPEi (FigS5L) challenged mitosis was also noticed, without further impact on mitotic timing (Figure S5J).

Cytoplasmic actin contributes to MT dynamics, mitotic spindle assembly and chromosome segregation (Mogessie & Schuh, 2017; Colin *et al*, 2018; Kita *et al*, 2019; Plessner *et al*, 2019). In our 3D experiments, actin depolymerization resulted in decreased MT polymerization rates and reduced chromosome mis-segregations. Notwithstanding, it was important to reinforce the possibility that cortical actin could be a critical factor in constrained conditions. We therefore tested the consequences of actin depolymerization in 2D aRGs, which are not subjected to constraints imposed by neighbors during mitosis. Interestingly, even if LatB treatment led to a decrease in cortical actin levels (Figure S6A), it did not impact the number or length of MTs polymerized from the poles (Figures S6B-C), nor the rate of chromosome mis-segregation (Figures S6D-E).

To further investigate the role of cortical actin as a sensor of biomechanical compression in mitosis, we perturbed branched actin nucleation *via* an acute treatment with the Actin-Related Protein 2/3 (Arp2/3) complex inhibitor CK666 (at 100μM for 1h-2.5h) (Hetrick *et al*, 2012). As expected from a decrease in branched actin polymerization activity by Arp2/3 complex inhibition, a switch from lamellipodia to filopodia structures (Suraneni *et al*, 2012) at the interface between 3D aRG cultures and the culture support was noticed (Figure 5G). Furthermore, we observed a delocalization of ArpC5, a subunit of the Arp2/3 complex from the plasma membrane to the cytoplasm (Figures 5H-I). Strikingly, both the MT nucleation capacity from the spindle poles (Figure 5J-K) and the rate of chromosome mis-segregation (Figure 5L), without obvious alterations in mitotic timing (Figure S6F), were significantly decreased in 3D aRG cells after Arp2/3 inhibition. Moreover, no such reduction in chromosome mis-segregation frequency was observed after CK666 treatment of 2D aRG cultures (Figures S6G-H).

Our results show that perturbing cortical cell properties —either by shortening actin filaments or decreasing the polymerization of branched actin— is sufficient to concurrently alter polar MT polymerization and the frequency of chromosome segregation errors exclusively in aRG cells under constraint.

### Cortical actin acts upstream of MCAK to tune polar MT polymerization and modulate chromosome segregation errors

We next determined whether cortical actin and MCAK depolymerase activity act in the same pathway. To do so, we combined UMK57 and LatB treatments in 3D aRG cultures. Interestingly, there were no differences in polar MT polymerization rates (Figures 5M-N) nor chromosome mis-segregation frequency (Figure 5O) in 3D aRG cells treated with UMK57 in the presence or absence of LatB-induced actin depolymerization. Worth mentioning that although the combination of UMK57 and MPS1i treatments caused a delay in anaphase onset (Figure S3I), which was further enhanced in the presence of LatB (Figure S6I). However, there was no obvious correlation with the absence of chromosome segregation errors (Figure 5O).

Together, these results advocate that biomechanical constraints faced by aRG mitosis are sensed by cortical actin and conveyed to MCAK depolymerase to balance polar MT polymerization rates and chromosome segregation accuracy.

### Releasing biomechanical constraints *in vivo* modulates astral MT polymerization and governs mitotic fidelity

Our results show that cortical actin levels link polar MT polymerization and chromosome mis-segregation rates in mitotic aRG cells under constrained conditions imposed by neighboring cells. We next tested whether releasing biomechanical constraints *in vivo* would alleviate the levels of chromosome mis-segregation observed at the early stage - E13.5, when aRG crowding is elevated. Incubation of cerebral cortex explants with LatB was sufficient to disrupt the ventricular lining in a focal manner (Figure S7A), validating the release of membrane tension in brain aRG cells. Similar to the experiments performed *in vitro*, the length and number of polar MTs after depolymerization/repolymerization assays were decreased (Figures S7B-D), which directly impacted the frequency of chromosome mis-segregation (Figures S7E-F). We next incubated cerebral cortex explants with CK666 as before. This treatment was sufficient to delocalize Arp2/3 complexes from the cortical membrane of aRG apical processes (Figure 6A). Further, in mitotic aRG cells cortical actin appeared wavy, as compared to control (Figure 6B). Importantly, this treatment was sufficient to trigger a drop in the number and length of EB1 comets polymerized from the spindle poles (Figure 6C-D) and a decrease in the rate of chromosome mis-segregation events (Figure 6E-F).

**Figure 6:**
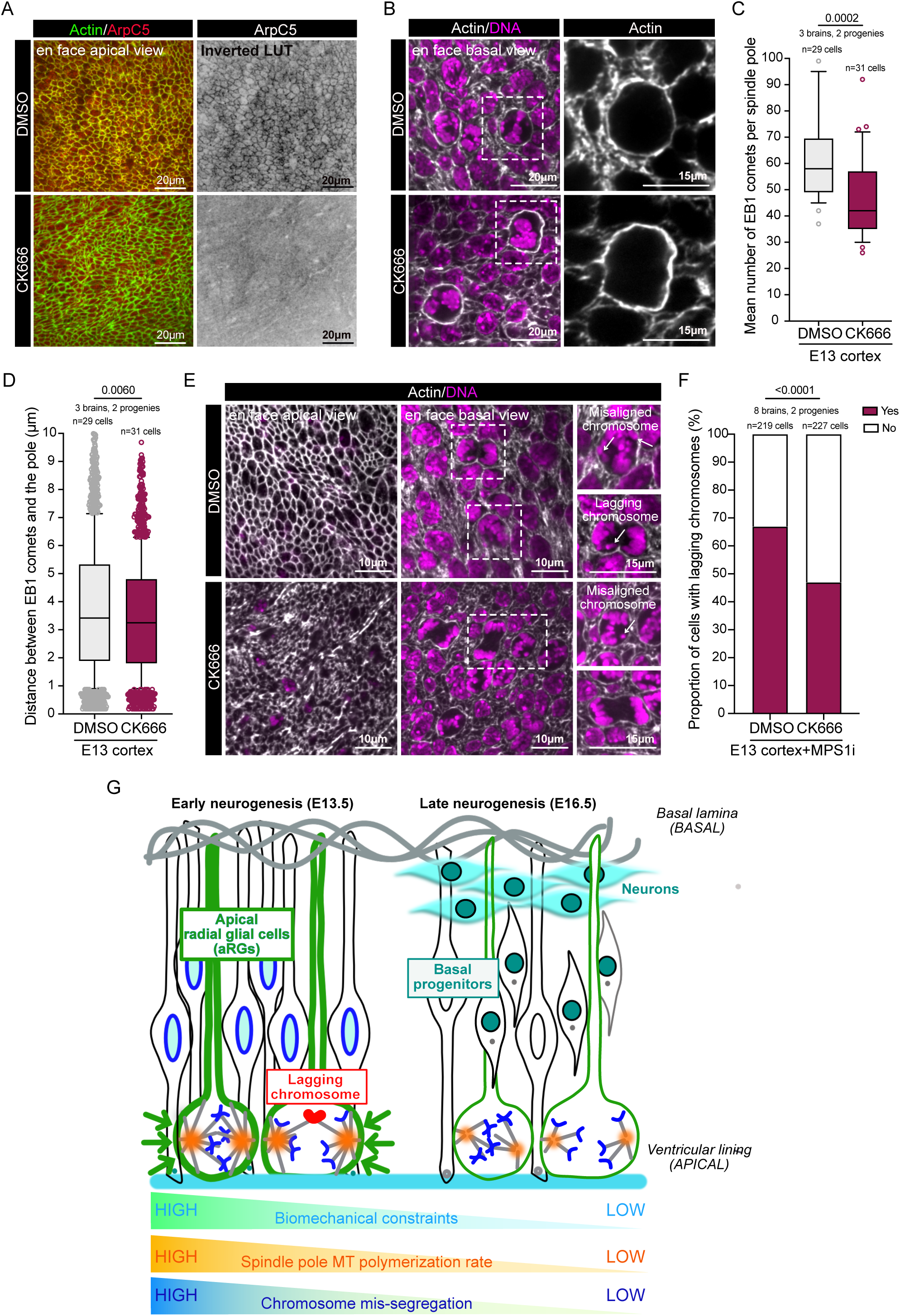
Released cortical tensions in the embryonic cerebral cortex influences mitotic accuracy through modulation of spindle pole MT polymerization. (A) Representative pictures of E13.5 cortical explants labelled with antibodies that recognize ArpC5 (red) and actin (green) as indicated. (B) Representative *en face* views of embryonic cortical explants treated with either DMSO (upper panels) or CK666 (bottom panels) showing actin (green) and DNA (magenta). (C-D) Box and whiskers plot showing the number of EB1 comets polymerized per pole (C) and the distance of individual EB1 comets from the spindle pole (D) during 3 min of MT repolymerization as indicated. (E) Representative images of *en face* views of embryonic cortical explants treated as indicated, showing actin (grey) and DNA (magenta). Left panels show *en face* apical views of the ventricular zone and middle panels basal views of aRG cells in mitosis. Right panels display corresponding high magnifications of aRG cells in prometaphase (top panels for each condition) and anaphase (bottom panels for each condition) highlighted with dotted rectangles in lower magnification basal views. The white arrows point either to a mis-aligned chromosome in metaphase or a lagging chromosome in anaphase. (F) Bar graph showing the proportion of cells with lagging chromosomes as indicated. (G) Model illustrating the impact of aRG cell density on spindle assembly and MT polymerization rates influencing chromosome segregation during brain development. Data were submitted to normality and lognormality tests for the choice of appropriate statistical test. Statistical significance was assessed with Mann-Whitney tests (C and D) and Fisher’s exact test in (F).

In conclusion, the cortical stiffness of mitotic aRG cells, conveyed by branched actin polymerization at the plasma membrane, acts as a modulator of MT nucleation at the spindle poles and consequently in chromosome segregation accuracy in the developing brain.

## Discussion

Cell division requires the orchestration of key events such as cell rounding, chromosome condensation and mitotic spindle assembly. Such a complex event relies on cell-autonomous and non-cell autonomous factors that sense and coordinate the activity of many mitotic players to achieve a common goal -generating two genetically identical euploid daughter cells. Although aneuploidy—the gain or loss of whole chromosomes (Santaguida, 2024; Maiato & Silva, 2023)—is known to depend on multiple factors and is often described as tissue context-dependent (Ben-David & Amon, 2020), the precise definition of this context remains poorly understood. Here, we show that the mechanical forces at play in the neocortex during neurogenesis have a critical impact on chromosome segregation fidelity in the neural progenitor population. At early stages of neurogenesis, when aRG cells undergo mitosis in a crowded environment, these contain high levels of cortical actin. This results in increased cortical stiffness, which modulates the rate of MT polymerization at the poles of the mitotic spindle. In particular, we show that high polar MT polymerization rates imposed by high cell density result in a higher probability of chromosome mis-segregation (Figure 6G). These findings link cell crowding with cortical actin stiffness and MTs from the mitotic spindle. More importantly, they show that cell crowding ultimately impact chromosome segregation fidelity. The brain is a complex organ where multiple cell lineages are generated within a relatively short period of time and, importantly, cell division in the cerebral cortex remains largely restricted to embryonic development. aRG cells divide extensively in the ventricular zone at early stages of neurogenesis, generating a crowded environment at the apical side of the neuroepithelium. We and others have previously reported that astral MT-enriched spindles, which are particularly abundant at early stages of neurogenesis in the aRG population (Mora-Bermúdez *et al*, 2014), are prone to generate mitotic errors (Vargas-Hurtado *et al*, 2019; Marthiens & Basto, 2020). Our results now show that it is the level of mechanical stress faced by mitotic aRG cells that favors the formation of astral MT-enriched spindles by promoting the polymerization of MTs from spindle poles. How MTs polymerized from the poles can fuel erroneous chromosome attachments and consequent mis-segregation remains to be investigated. Likely, abnormal chromosome-MT attachments that escape the SAC surveillance, i.e., merotelic attachments (Cimini *et al*, 2001) may be promoted in the presence of more abundant and/or dynamic population of astral MTs probing the cellular space to establish connections with kinetochores.

Few studies have examined the role of tissue environment in influencing mitotic fidelity. It has recently been shown that epithelial cells show increased chromosome segregation errors after polarity loss outside of their native environment and in an integrin-dependent manner (Knouse *et al*, 2018). Our findings support the same view that tissue properties influence chromosome segregation fidelity, but they suggest instead that within the developing mouse brain, a major contribution from cell crowding and cortical actin has to be considered.

It is tempting to speculate that during brain development, the reason why aRG cells are particularly prone to mitotic errors in pathological conditions relies on their unique mechano-sensitivity at early neurogenic stages. Our findings put forward the view that brain biomechanical properties have to be considered when defining cell division vulnerability and chromosome instability in adverse situations.

## Acknowledgements

We acknowledge the Cell and Tissue Imaging platform (PICT-IBiSA), Institut Curie, member of the national infrastructure France-BioImaging (https://ror.org/01y7vt929) supported by the French National Research Agency (ANR-24-INBS-0005 FBI BIOGEN) for help and support with image acquisition and analysis. We acknowledge S. Fre, A. M. Lennon and J.L. Maitre (Institut Curie, Paris) for providing mT/mG, LifeAct-GFP and mCherry-H2B mouse lines, respectively. We thank A. Terrizzano and A. Simon (Institut Curie) for invaluable help with mouse sample preparations and colony management. We thank O. Goundiam for helpful comments on statistical analysis. We thank A. Baffet (Institut Curie), G. Cappello (Grenoble-Alpes University, France), J.B. Manneville (Paris-Cite University, France), S. Passemard (HU R. Debre, Paris, France), N. Pavin (University of Zagreb, Croatia), M. Piel (Institut Curie), D. Matic-Vignjevic (Institut Curie) and M-E. Terret (CIRB, College-de-France, Paris) for stimulating discussions. We thank all past and present members of the Basto lab for helpful discussions and critical comments on the manuscript, and especially F. Edwards, S. Gemble, O. Goundiam, F. Scotto di Carlo, A. Simon and A. Terrizzano. V.M. is a permanent INSERM researcher. E.L. was supported by grants CEECIND/00654/2020 and PTDC/MED-OUT/2747/2020 from Fundação para a Ciência e a Tecnologia (FCT).This work received support from a collaborative grant to V.M. and C.V. funded by the Labex Cell(n)scale from ANR-11-LABX-0038 and ANR-10-IDEX-0001-02, a MiCMac ANR (ANR22-CE16-0008) to R.B. and the ERC consolidator grant to R.B (725907 ERC-2016-COG).

## Author contributions

V.M. and R.B. conceived the project. V.M wrote the manuscript. V.M. did most of the experiments and data analysis presented here with help from C.B, L.J. M.M. and E.L. D.K. provided help and expertise with the laser cut experiments, O.Z. and L.P. with the dextran experiments and Arp2/3 complex perturbation assays, respectively. A.S.M. generated the macros used to quantify cell volumes and extract EB1 comet growth parameters. C.P, T.P and C.V. designed and generated the PDMS support with 3D grooves of different dimensions and performed in-mold patterning as tools to confine aRG cells *in vitro*. E.L provided the UMK57 agonist and expertise to perturb MCAK activity. V.M and R.B. obtained the funding used to develop this project. All authors read and commented on the manuscript.

## Materials and Methods

### Animal Husbandry and experimental procedures

Concerning animal care, we followed the European and French National Regulation for the Protection of Vertebrate Animals used for Experimental and other Scientific Purposes (Directive 2010/63; French Decree 2013-118). The project was authorized and benefited from guidance of the Animal Welfare Body, Research Centre, Institut Curie. All mice were kept in the Institut Curie Specific Pathogen Free (SPF) animal facility for breeding. C57Bl6/N pregnant females were purchased from Charles River Laboratories (France). The transgenic mT/mG mouse line (stock number 007576, The Jackson Laboratory) (Muzumdar *et al*, 2007), LifeAct-GFP mouse line (Riedl *et al*, 2008), and Rosa26-mCherry-H2B mouse line (Abe *et al*, 2011) were inbred in our animal facility. For all pregnant females, the day of mating was referred to 0.5 day of gestation (E0.5). Pregnant females were anaesthetized by inhalation of isoflurane (Baxter, DDG9621) before being sacrificed by cervical dislocation. Embryos were collected after cesarean section.

### Preparation of cortical explants, 3D or 2D apical radial glia (aRG) primary cultures

Dorsal telencephalons (referred after as cerebral cortices or cortical explants) of wild type C57Bl6/N mice were dissected at embryonic day 13.5 (E13.5) or 16.5 (E16.5) of neurogenesis. Preparations of mouse embryonic brain tissues for *en face view* analysis were described in detail in (Rujano *et al*, 2015). After dissection, intact cortical explants were incubated for 15 min at 37°C under 5% CO_2_ in proliferation medium before any other procedures. Proliferation medium contains DMEM/HamF12 medium (Gibco, 42400-028), supplemented with glucose (Sigma, G7021, 2.9g/L), sodium bicarbonate (Sigma, S5761, 1.2g/L), B27 supplement w/o vitamin A (Gibco, 12587-010; 2%), Fibroblast Growth Factor (Human FGF basic (FGF-2) 146a.a, Peprotech, 100-18C, 20 ng/ml), Epidermal Growth Factor (Recombinant murine EGF, Peprotech, 31509, 20 ng/ml) and antibiotic-antimycotic 1% (Gibco, 15240-062).

To establish primary cultures of apical radial glia (aRG), we have adapted the protocol described in (Sun *et al*, 2011). Cortical explants were dissociated into a single cell suspension by a combination of enzymatic treatment -incubation for 5 min at 37°C with accutase (Sigma, A6964)- and mechanical dissociation of mouse E13.5 brains. To establish 3D aRG cultures, cells were maintained as 3D floating aggregates -also called neurospheres-in non-adherent flasks at 37°C under 5% CO_2_. Cells were cultured for one week in proliferation medium to enable maximal enrichment for Sox2-positive apical radial glia (aRG) before any experimental procedure (Vargas-Hurtado *et al*, 2019). They were kept in culture for 4 weeks.

Before using drugs and immunofluorescent approaches or time-lapse microscopy, 3D aRG cells were distributed on poly-D-Lysine coated glass/plastic supports for 2H. For live imaging, neurospheres were plated on PDL-coated glass bottom dishes (Greiner Bio-one, 627870) 2H before the start of the experiment. To establish 2D aRG cultures for perturbation assays, cells were dissociated the day before from neurospheres and plated on plastic/glass supports pre-coated sequentially for 2H at RT with poly-D-lysine hydrobromide (PDL, Sigma, P6407, 2μg/cm^2^) and fibronectin (Sigma, F1141, 1μg/cm^2^).

Transfections were performed with the Amaxa mouse NSC nucleofector kit (Lonza, VPG-1004), according to the manufacturer’s instructions. pGFP-EB1 was a gift from Lynne Cassimeris (RRID:Addgene_17234, http://n2t.net/addgene:17234) (Piehl & Cassimeris, 2003).

### Imposed mechanical stress on 2D aRG cells

To achieve bi-functionalized PDMS trenches, we adapted a technique previously developed by (Yamada *et al*, 2016) named in-mold patterning (IMP). Briefly, this technique uses a Polydimethylsiloxane (PDMS) stamp of inverted trench polarity and a PDMS inker. The inker is incubated with 10% sodium dodecyl sulfate (SDS) solution for 15min, rinsed with water and further incubated with a 100μg/ml fibronectin (FN) solution for at least 1h. The stamp is exposed to an oxygen plasma, silanized using (Tridecafluoro-1,1,2,2-tetrahydrooctyl) trichlorosilane (ABCR, AB111444) for 30min, and incubated with 10% SDS solution for 30min before being dried with pressurized nitrogen. Lastly, the stamp is pressed for 3min on the inker using a 30g metal weight (the FN solution is pipetted just before, preventing the inker from drying before the stamping process) and removed. The inked stamp is further spin-coated with fresh PDMS at 500rpm for 10s, then 3000rpm for 30s (for a final theoretical thickness of about 75μm). A plasma activated glass coverslip is then placed on top of the PDMS layer as a physical support which, after reticulation at 70°C overnight, will stick to the IMP. After peeling off from the stamp, we obtain an easy-to-handle structured substrate with 10µm deep trenches functionalized with FN at their bottom. The sample is next incubated for 1h in a 4µg/mL Ncad-Fc solution (BD, 1388-NC), deposited in the form of a drop on top of it, to functionalize the walls and the upper surface of the trenches. Importantly, we noticed that the Ncad-Fc did not cover the FN. At last, to remove Ncad-Fc from the surface of the IMP substrate and avoid cells to adhere elsewhere than inside the trenches, we used the stamp-off method originally proposed by (Desai *et al*, 2011), consisting in placing in conformal contact a flat, plasma activated PDMS layer, over the IMP substrate for 30s. The top Ncad-Fc is then transferred to this PDMS stamp-off object.

To impose mechanical stress on 2D aRG and perform live imaging, aRG cells dissociated from neurospheres were cultured on a PDL/fibronectin substrate and incubated for 1h (fixed samples) or 2.5h (live imaging) in the presence of a solution of Dextran (Sigma, D5376) prepared at a final concentration of 80mg/mL in the proliferation medium. Indeed, it has been proven (Dolega *et al*, 2021) that the osmotic pressure Π due to the osmolyte directly results in a solid stress σ of identical magnitude (σ = Π), under two conditions: 1) in the limit where the compressibility of the fluid is much lower than that of the solid phase. This condition is met here, as the bulk modulus of water is more than thousand times larger than that of cells (Monnier *et al*, 2016), 2) when the osmolyte cannot be internalized by the cells, which is the case for dextran polymers of large molecular weight (Zlotek-Zlotkiewicz *et al*, 2015). Dextran of large molecular weight are long polymers. Thus, their osmotic pressure cannot simply be computed using van ’t Hoff approximation, which is valid only for very diluted solutions. To determine the precise dilution required to obtain the desired pressure, we used the calibration curves published previously by (Vink, 1971; Bonnet-Gonnet *et al*, 1994).

Dissociated aRG cells were coated on a substrate of Ncadherin-Fc chimera (NcadFc, BD, 1388-NC) to mimic N-cadherin-dependent cell-cell contacts (Gavard *et al*, 2004). Ncad-Fc substrate was prepared by incubating PDL-coated supports with a solution of Ncad-Fc at 10μg/mL (BD, 1388-NC) prepared in Leibowitz’s L-15 medium (ThermoFisher Scientific, 11415064).

### Microtubule depolymerization-repolymerization experiments and drug-based assays

For microtubule (MT) depolymerization-repolymerization assays, cortical explants or aRG cultures were incubated for 1h at 4°C in proliferation medium to depolymerize MTs. Explants/cells were then incubated between 1.5min and 3min at 37°C for MT repolymerization in the same medium before fixation for 2min in −20°C precooled methanol and immunofluorescent staining.

For drug treatments, the appropriate concentrations were predetermined in adherent aRG cultures. Chromosome misalignment and mis-segregation events were promoted using inhibitors of CENPE motor activity (CENPEi, GSK923295, Selleckchem, S7090) at 400nM and MPS1 activity (MPS1i, MPI-0479605, Selleckchem, S7488) at 25nM on 2D or 3D aRG cultures or 100nM on E13.5 cerebral cortex explants. Actin filament polymerization was prevented using Latrunculin B (Millipore, 428020) at 50nM. Branched actin polymerization was prevented using CK666 (Tocris Bioscience, 3950) at 100μM on 2D or 3D aRG cultures and 200-400μM on E13.5 cerebral cortex explants. MT dynamics was perturbed after treatment with the MCAK agonist UMK57 at 1μM (Barroso-Vilares *et al*, 2020). All the drugs were diluted in dimethyl sulfoxide anhydrous (DMSO, Sigma, 276855). Explants/cells incubated with DMSO alone, diluted at equivalent concentrations (V/V), were used as negative controls. Explants /cells were incubated for 1h with the drugs before fixation and immunofluorescent staining or for 2.5h during live imaging.

### Immunofluorescence approaches

Cells/explants were fixed using different methods depending on the primary antibodies used. Cells or explants were either fixed for 15 or 30min, respectively, in paraformaldehyde (PFA), diluted at 4% in PBS at room temperature, or fixed for 2min in −20°C precooled methanol. Samples were further incubated for 1h in blocking solution (PBS, BSA 3%, Triton X-100 0.3%, Sodium azide 0.02%), followed by sequential overnight primary and secondary antibody incubations in the same buffer. Washes were performed with the blocking solution after each incubation. Cortical explants were mounted with the apical surface facing the glass coverslips for further analysis of apically localized mitotic progenitors, as described in (Rujano *et al*, 2015).

The following primary antibodies were used: mouse IgG1 anti-α-tubulin, dilution 1/500 (clone DM1α, Sigma, T6199, RRID:AB_477583), human anti-centromere, dilution 1/100 (Antibodies Incorporated, 15-234-0001, RRID:AB_2687472), mouse IgG1 anti-N-cadherin, dilution 1/500 (BD Transduction Laboratories, 610921, RRID:AB_398236), rabbit anti-phospho-ERM, dilution 1/500 (Cell Signaling Technology, 3141, RRID:AB_330232), goat anti-Sox2, dilution 1/500 (R&D, AF2018, RRID:AB_355110) and mouse IgG2a anti-p16Arc, dilution 1/500 (ARPC5 subunit, Synaptic System Antibodies, 305011, RRID:AB_887896), mouse IgG1 anti-EB1, dilution 1/500 (Cell Signaling, 2164S, RRID:AB_2141765) and rabbit anti-phospho-histone 3 (Ser10), dilution 1/500 (Sigma, H0412, RRID:AB_477043). All secondary antibodies used were highly cross-absorbed: goat anti-rabbit Alexa Fluor 488 (Thermo Fisher Scientific, A-11034, RRID:AB_2576217), goat anti-rabbit Alexa Fluor 546 (Thermo Fisher Scientific, A-11035, RRID:AB_2534093), goat anti-rabbit Alexa Fluor 647 (Thermo Fisher Scientific, A-21245, RRID:AB_2535813), Cy5 AffiniPure Donkey Anti-Rabbit (Jackson Immunoresearch, 709-175-149, RRID:AB_2340539), goat anti-mouse IgG1 Alexa Fluor 488 (Thermo Fisher Scientific, A-21121, RRID:AB_2535764), goat anti-mouse IgG2a Alexa Fluor 546 (Thermo Fisher Scientific, A-21133, RRID:AB_2535772), donkey anti-goat IgG Alexa Fluor 488 (Thermo Fisher Scientific, A-32814, RRID:AB_2762838), Cy3 AffiniPure Donkey Anti-Human (Jackson Immunoresearch, 709-165-149, RRID:AB_2340535). For actin labeling, Alexa Fluor 488 Phalloidin (Thermo Fisher Scientific, A-12379), Alexa Fluor Plus 555 Phalloidin (Thermo Fisher Scientific, A-30106), Alexa Fluor Plus 647 Phalloidin (Thermo Fisher Scientific, A-30107) were used at 1/500 dilution. 4’,6-diamidino-2-phenylindole (DAPI, Sigma, D1306) was used to label DNA at a concentration of 3μg/ml and incubated overnight at RT at the same time than secondary antibodies. EverBrite Mounting medium (Biotium, 23001) was used for mounting.

### Cortical membrane factor recruitment analysis

For *in vivo* analysis, actin and phospho-ERM (p-ERM) protein levels at the cortical membrane were related to N-cadherin protein levels, which do not change between early and late neurogenesis. A 5 pixel width line scan fluorescence intensity analysis was performed for each marker along the plasma membrane identified with the N-cadherin staining. The mean intensity value was evaluated for each marker of interest and related to the mean intensity value of N-cadherin at the plasma membrane.

A similar 5 pixels width line scan fluorescence intensity analysis was performed to evaluate actin fluorescence intensity (FI) levels at the plasma membrane after drug treatment in 3D or 2D aRG primary cultures. The mean intensity value was plotted for each mitosis.

### aRG mitosis morphometrics analysis

To evaluate aRG mitotic cell volumes in complex 3D environment and delineate cell contour, we took advantage of the cytoplasmic dispersion of the stem cell transcription factor Sox2 after nuclear envelope breakdown. For volume computation, Cellpose algorithm (CellPose2, RRID:SCR_021716, http://www.cellpose.org) (Pachitariu, 2022) was launched on the Sox2 channel on each slice separately in batch (Figure S1D). For each stack, a Fiji macro was developed where the user choses the cells of interest by drawing a rectangle around them. The largest label obtained from Cellpose segmentation on each slice is stored in the ROI Manager. The user can manually refine the segmentation if needed, by modifying segmentation on some slices or adding missing regions (especially for out-of-focus slices where the segmentation was less accurate). The volume is then computed using the sum of area multiply by the interval between optical slices (Z step) (Figure S1D). The dedicated macro can be found in https://github.com/Anne-SophieMACE/MitoticCellVolume_EB1cometAnalysis. 3D reconstructions of mitotic cell volumes displayed in Figure 1F were performed using the Imaris software (RRID:SCR_007370, http://www.bitplane.com/imaris/imaris).

### Cortical membrane tension measurement by laser ablation

Embryonic cortical explants prepared from LifeAct-GFP (cortical actin endogenously labelled with GFP) expressing transgenic mice progeny were placed for *en face view* acquisitions with the apical surface down facing the glass bottom tissue culture dishes (Fluorodish, FD35-100, WPI) in 80μL of 37°C prewarmed medium. Explants were maintained in place during the procedure with a slice anchor (Slice Anchor Kit for RC-22, SHD-22KIT, 64-0263, Warner Instruments) to minimize sample drift, surrounded with 10S Voltalef oil (VWR Chemicals, 24627.188) to favour medium oxygenation and prevent evaporation during the whole procedure. 3D aRG cultures prepared from mT/mG (membrane-targeted tandem dimer Tomato (mT)) expressing transgenic mice progeny were plated on PDL-precoated glass bottom four compartment cell culture dishes (Greiner Bio-one, 627870). Tissues/cells were imaged using a two-photon laser-scanning microscope (LSM880, Carl Zeiss) in single photon mode (488 or 561 line), using a 63x/NA 1.4 OIL DICII PL APO objective. The cortical membrane was ablated using a Ti:Sapphire laser (Mai Tai DeepSee, Spectra Physics) set at 800nm. The laser power was optimized to efficiently ablate the cortical membrane without creating cavitation or cell damage. A single Z-section was considered at the equator of each mitotic cell for laser cut and acquisition. Images were acquired with a time interval δt=100 ms.

Image quality was improved using the Fiji software gaussian blur filter to improve signal-to-noise and increase cortical membrane localization accuracy. Cortical membrane tension was evaluated by measuring the initial velocity recoil post-ablation, corresponding to ratio of the size of the wound in mm over the time elapsed between laser cut and first image recording (based on measured δt).

### Image acquisitions on fixed samples and time-lapse recording

Images were acquired on a spinning disk wide version microscope (Gataca Systems, France). Based on a CSU-W1 (Yokogawa, Japan), the spinning head was mounted on an inverted Eclipse Ti2 (Nikon) microscope equipped with a motorized XY Stage (Nikon, Japan). Images were acquired through 100x 1.4NA Plan-Apo (fixed samples) or 40X 1.3NA Plan-Apo (live imaging) objectives with a sCMOS camera (Prime95B, Photometrics, USA). Optical sectioning was achieved using a piezo stage (Nano-z series, Mad City Lab, USA). Gataca Systems’ laser bench was equipped with 405, 491 and 561 nm laser diodes, delivering 150 mW each, coupled to the spinning disk head through a single mode fiber. Multi-dimensional acquisitions were performed using MetaMorph 7.10.1 software (MetaMorph Microscopy Automation and Image Analysis Software, RRID:SCR_002368, http://www.moleculardevices.com/Products/Software/Meta-Imaging-Series/MetaMorph.html). Image analyses were performed using free online Fiji software (Image J2 software, RRID:SCR_003070 https://imagej.net/software/fiji/downloads). For time-lapse recording and scoring of chromosome segregation errors, mCherry-H2B expressing aRG cells in 2D or 3D were plated on the appropriate substrates in glass-bottom four compartment cell culture dishes (627870, Greiner BioOne). Images were acquired every 2min with 1μm z intervals for a maximum of 20z stacks per position during 2.5 to 3h after adding the drugs. Mitotic timing was calculated as the time elapsed between nuclear envelope breakdown (NEBD) and anaphase onset. A mean mitotic timing between 20 and 30min in the control aRG population was considered as a quality control criterion to select the primary cultures to be used for analysis. Any control aRG population displaying more than 30min mitotic timing was discarded from the analysis. The frequency of chromosome segregation errors in the population was scored as the number of chromosomes lagging at t=4min after anaphase onset (second time point recorded after chromatid separations). Data were plotted using GraphPad Prism (version 10 for Mac, GraphPad software, RRID:SCR_002798, www.graphpad.com) and statistical analysis performed with the same software. Figures were mounted using Affinity Designer software (Affinity Designer, RRID:SCR_016952, https://affinity.serif.com/en-us/designer/).

### MT dynamics analysis after live imaging or MT depolymerization-repolymerization assays

For tracking of EB1-positive comets in timelapse movies of mitotic cells, the spindle pole with the highest MT polymerization activity was chosen. Acquisitions were performed at a single z section during 1min every 1sec in prometaphase with a 100X objective mounted on the spinning disk wide version microscope (Gataca Systems, France), described in the image acquisitions section. We chose for analysis spindle poles stable in z and positioned in *en face* views so that tracked comets were not lost because of z axis movement. For each movie, we chose randomly three time points during the one-minute recording. We tracked each EB1 positive comet arising from one single pole until its disappearance. We evaluated the number of comets arising from the pole at a given time point (MT polymerization rate) and the maximum distance they travelled before disappearing (evaluation of the MT polymerization persistence).

A semi-automated Fiji macro was developed to compute EB1-positive comet number, length and distance travelled from the pole on fixed samples after MT depolymerization-repolymerization assays. The analysis was performed on a 0.4mm maximum z-projection (3 z sections acquired at intervals of 0.2mm). The user is asked first to draw the pole and the area where comets should be detected. The comets are detected as local maxima using the Find Maxima Plugin in Fiji, with the prominence parameter chosen by the user to detect most comets; manual editing of comets is possible by the user. Using the ROI of the pole, the distance from each comet to the pole is easily computed with the Fiji Plugin “Distance Transform 3D”. For measuring comet length, we considered the fact that all comets propagate outward from the pole and that the length can be determined by analysing the line passing through the pole and the comet centre. Along this line, a Gaussian fit centred on the comet’s position is performed; the length of the comet is then defined as the FWHM (full width at half maximum), approximated by 2.355 sigma. Aberrant values or bad fit are excluded, and the user can again manually check and adjust comets whose size were not well approximated (for instance, comets rotated relative to the pole/comet line, or those too close to another comet). The dedicated macro can be found in https://github.com/Anne-SophieMACE/MitoticCellVolume_EB1cometAnalysis.

## Bibliography

Abe T, Kiyonari H, Shioi G, Inoue KI, Nakao K, Aizawa S & Fujimori T (2011) Establishment of conditional reporter mouse lines at ROSA26 locus for live cell imaging. Genesis 49: 579–590

Arai Y & Taverna E (2017) Neural progenitor cell polarity and cortical development. Front Cell Neurosci 11: 1–11

Barroso-Vilares M, Macedo JC, Reis M, Warren JD, Compton D & Logarinho E (2020) Small-molecule inhibition of aging-associated chromosomal instability delays cellular senescence. EMBO Rep 21: e49248

Ben-David U & Amon A (2020) Context is everything: aneuploidy in cancer. Nat Rev Genet 21: 44–62

Bonnet-Gonnet C, Belloni L & Cabane B (1994) Osmotic pressure of latex dispersions. Langmuir 10: 4012–4021

Cai X, Evrony GD, Lehmann HS, Elhosary PC, Mehta BK, Poduri A & Walsh CA (2014) Single-Cell, Genome-wide Sequencing Identifies Clonal Somatic Copy-Number Variation in the Human Brain. Cell Rep 8: 1280–1289

Carreno S, Kouranti I, Glusman ES, Fuller MT, Echard A & Payre F (2008) Moesin and its activating kinase Slik are required for cortical stability and microtubule organization in mitotic cells. Journal of Cell Biology 180: 739–746

Casas Gimeno G & Paridaen JTML (2022) The Symmetry of Neural Stem Cell and Progenitor Divisions in the Vertebrate Brain. Front Cell Dev Biol 10: 1–21

Chugh P, Clark AG, Smith MB, Cassani DAD, Dierkes K, Ragab A, Roux PP, Charras G, Salbreux G & Paluch EK (2017) Actin cortex architecture regulates cell surface tension. Nat Cell Biol 19: 689–697

Cimini D, Howell B, Maddox P, Khodjakov A, Degrassi F & Salmon ED (2001) Merotelic kinetochore orientation is a major mechanism of aneuploidy in mitotic mammalian tissue cells. Journal of Cell Biology 152: 517–527

Colin A, Singaravelu P, Théry M, Blanchoin L & Gueroui Z (2018) Actin-Network Architecture Regulates Microtubule Dynamics. Current Biology 28: 2647–2656.e4

Desai RA, Khan MK, Gopal SB & Chen CS (2011) Subcellular spatial segregation of integrin subtypes by patterned multicomponent surfaces †.

Dolega ME, Monnier S, Brunel B, Joanny JF, Recho P & Cappello G (2021) Extra-cellular matrix in multicellular aggregates acts as a pressure sensor controlling cell proliferation and motility. Elife 10: 1–33

Farcy S, Hachour H, Bahi-Buisson N & Passemard S (2023) Genetic Primary Microcephalies: When Centrosome Dysfunction Dictates Brain and Body Size. Cells 12: 1–34

Farhadifar R, Röper JC, Aigouy B, Eaton S & Jülicher F (2007) The Influence of Cell Mechanics, Cell-Cell Interactions, and Proliferation on Epithelial Packing. Current Biology 17: 2095–2104

Gavard J, Marthiens V, Monnet C, Lambert M & Mège RM (2004) N-cadherin Activation Substitutes for the Cell Contact Control in Cell Cycle Arrest and Myogenic Differentiation. Journal of Biological Chemistry 279: 36795–36802

Gogendeau D, Siudeja K, Gambarotto D, Pennetier C, Bardin AJ & Basto R (2015) Aneuploidy causes premature differentiation of neural and intestinal stem cells. Nat Commun 6: 1–15

Gudimchuk N, Vitre B, Kim Y, Kiyatkin A, Cleveland DW, Ataullakhanov FI & Grishchuk EL (2013) Kinetochore kinesin CENP-E is a processive bi-directional tracker of dynamic microtubule tips HHS Public Access. Nat Cell Biol 15: 1079–1088

van Heesbeen RGHP, Raaijmakers JA, Tanenbaum ME, Halim VA, Lelieveld D, Lieftink C, Heck AJR, Egan DA & Medema RH (2017) Aurora A, MCAK, and Kif18b promote Eg5-independent spindle formation. Chromosoma 126: 473–486

Hetrick B, Suk Han M & Nolen B (2012) Small Molecules CK-666 and CK-869 Block an Activating Conformational Change to Inhibit Arp2/3 Complex. Biophys J 102: 46a

Jayaraman D, Bae B Il & Walsh CA (2018) The genetics of primary microcephaly. Annu Rev Genomics Hum Genet 19: 177–200

Kaindl AM, Passemard S, Kumar P, Kraemer N, Issa L, Zwirner A, Gerard B, Verloes A, Mani S & Gressens P (2010) Many roads lead to primary autosomal recessive microcephaly. Prog Neurobiol 90: 363–83

Kita AM, Swider ZT, Erofeev I, Halloran MC, Théry M & Saint-louis H (2019) Spindle – F-actin interactions in mitotic spindles in an intact vertebrate epithelium. Mol Biol Cell 30: 1645–1654

Knouse KA, Lopez KE, Bachofner M & Amon A (2018) Chromosome Segregation Fidelity in Epithelia Requires Tissue Architecture. Cell 175: 200–211.e13

Knouse KA, Wu J, Whittaker CA & Amon A (2014) Single cell sequencing reveals low levels of aneuploidy across mammalian tissues. Proc Natl Acad Sci U S A 111: 13409–13414

Krndija D, Marjou F El, Guirao B, Richon S, Leroy O, Bellaiche Y, Hannezo E & Vignjevic DM (2019) Active cell migration is critical for steady-state epithelial turnover in the gut. Science (1979) 365: 705–710

Kunda P, Pelling AE, Liu T & Baum B (2008) Moesin Controls Cortical Rigidity, Cell Rounding, and Spindle Morphogenesis during Mitosis. Current Biology 18: 91–101

Lancaster OM, LeBerre M, Dimitracopoulos A, Bonazzi D, Zlotek-Zlotkiewicz E, Picone R, Duke T, Piel M & Baum B (2013) Mitotic Rounding Alters Cell Geometry to Ensure Efficient Bipolar Spindle Formation. Dev Cell 25: 270–283

Lizarraga SB, Margossian SP, Harris MH, Campagna DR, Han AP, Blevins S, Mudbhary R, Barker JE, Walsh CA & Fleming MD (2010) Cdk5rap2 regulates centrosome function and chromosome segregation in neuronal progenitors. Development 137: 1907–1917

Machicoane M, de Frutos CA, Fink J, Rocancourt M, Lombardi Y, Gare S, Piel M & Echard A (2014) SLK-dependent activation of ERMs controls LGN-NuMA localization and spindle orientation. Journal of Cell Biology 205: 791–799

Maiato H & Silva S (2023) Double-checking chromosome segregation. Journal of Cell Biology 222 doi:10.1083/jcb.202301106 [PREPRINT]

Marjanović M, Sánchez-huertas C, Terré B, Gómez R, Scheel F, Pacheco S, Knobel PA, Martínez-marchal A, Palenzuela L, Wolfrum U, et al (2016) cep63 Reveals a Role for the Centrosome in Meiotic Recombination. Nat Commun

Marthiens V & Basto R (2020) Centrosomes: The good and the bad for brain development. Biol Cell: 1–20

Marthiens V, Rujano M a, Pennetier C, Tessier S, Paul-Gilloteaux P & Basto R (2013) Centrosome amplification causes microcephaly. Nat Cell Biol 15: 731–40

McConnell MJ, Lindberg MR, Brennand KJ, Piper JC, Voet T, Cowing-Zitron C, Shumilina S, Lasken RS, Vermeesch J, Hall IM, et al (2013) Mosaic copy number variation in human neurons. Science (1979) 342: 632–637

McHugh T & Welburn JPI (2023) Potent microtubule-depolymerizing activity of a mitotic Kif18b–MCAK–EB network. J Cell Sci 136

McIntyre RE, Lakshminarasimhan Chavali P, Ismail O, Carragher DM, Sanchez-Andrade G, Forment J V., Fu B, Del Castillo Velasco-Herrera M, Edwards A, van der Weyden L, et al (2012) Disruption of Mouse Cenpj, a Regulator of Centriole Biogenesis, Phenocopies Seckel Syndrome. PLoS Genet 8: 1–18

Mitchison T & Kirschner M (1984) Dynamic instability of microtubule growth. Nature 1984 312:5991 312: 237–242

Mogessie B & Schuh M (2017) Actin protects mammalian eggs against chromosome segregation errors. Science (1979) 357

Monnier S, Delarue M, Brunel B, Dolega ME, Delon A & Cappello G (2016) Effect of an osmotic stress on multicellular aggregates. Methods 94: 114–119

Mora-Bermúdez F, Matsuzaki F & Huttner WB (2014) Specific polar subpopulations of astral microtubules control spindle orientation and symmetric neural stem cell division. Elife 3

Muroyama A & Lechler T (2017) A transgenic toolkit for visualizing and perturbing microtubules in mice reveals unexpected functions for non-centrosomal arrays in epidermal morphogenesis. Elife 6: e29834

Muzumdar MD, Tasic B, Miyamichi K, Li L & Luo L (2007) A global double-fluorescent Cre reporter mouse. Genesis 45: 593–605

Naveed M, Kazmi SK, Amin M, Asif Z, Islam U, Shahid K & Tehreem S (2018) Comprehensive review on the molecular genetics of autosomal recessive primary microcephaly (MCPH). Genet Res (Camb) 100: 1–16

Pachitariu M, SC (2022) Cellpose 2.0: how to train your own model. Nat Methods 19: 1634–1641

Paridaen JT & Huttner WB (2014) Neurogenesis during development of the vertebrate central nervous system. EMBO Rep 15: 351–364

Piehl M & Cassimeris L (2003) Organization and Dynamics of Growing Microtubule Plus Ends during Early Mitosis. Mol Biol Cell 14: 916–925

Pilaz LJ, McMahon JJ, Miller EE, Lennox AL, Suzuki A, Salmon E & Silver DL (2016) Prolonged Mitosis of Neural Progenitors Alters Cell Fate in the Developing Brain. Neuron 89: 83–99

Plessner M, Knerr J & Grosse R (2019) Centrosomal Actin Assembly Is Required for Proper Mitotic Spindle Formation and Chromosome Congression. iScience 15: 274–281

Rehen SK, Yung YC, McCreight MP, Kaushal D, Yang AH, Almeida BSV, Kingsbury MA, Cabral KMS, McConnell MJ, Anliker B, et al (2005) Constitutional aneuploidy in the normal human brain. Journal of Neuroscience 25: 2176–2180

Riedl J, Crevenna AH, Kessenbrock K, Yu JH, Neukirchen D, Bista M, Bradke F, Jenne D, Holak TA, Werb Z, et al (2008) Lifeact: a versatile marker to visualize F-actin. Nat Methods 5: 605–607

Roostalu J, Thomas C, Cade NI, Kunzelmann S, Taylor IA & Surrey T (2020) The speed of GTP hydrolysis determines GTP cap size and controls microtubule stability. Elife 9: 1–22

Rujano MA, Basto R & Marthiens V (2015) New insights into centrosome imaging in Drosophila and mouse neuroepithelial tissues. Methods Cell Biol 129: 211–227

Rujano MA, Sanchez-Pulido L, Pennetier C, le Dez G & Basto R (2013) The microcephaly protein Asp regulates neuroepithelium morphogenesis by controlling the spatial distribution of myosin II. Nat Cell Biol 15: 1294–306

Santaguida S (2024) Two decades of chromosomal instability and aneuploidy. Chromosome Research 32: 1–3

Santaguida S, Tighe A, D’Alise AM, Taylor SS & Musacchio A (2010) Dissecting the role of MPS1 in chromosome biorientation and the spindle checkpoint through the small molecule inhibitor reversine. Journal of Cell Biology 190: 73–87

Shi L, Qalieh A, Lam MM, Keil JM & Kwan KY (2019) Robust elimination of genome-damaged cells safeguards against brain somatic aneuploidy following Knl1 deletion. Nat Commun 10: 1–14

Spector I, Shochet NR, Kashman Y & Groweiss A (1983) Latrunculins: Novel Marine Toxins That Disrupt Microfilament Organization in Cultured Cells. Science (1979) 219: 493–495

Stewart MP, Helenius J, Toyoda Y, Ramanathan SP, Muller DJ & Hyman AA (2011) Hydrostatic pressure and the actomyosin cortex drive mitotic cell rounding. Nature 469: 226–231

Stout JR, Yount AL, Powers JA, LeBlanc C, Ems-McClung SC & Walczak CE (2011) Kif18B interacts with EB1 and controls astral microtubule length during mitosis. Mol Biol Cell 22: 3070–3080

Sun T, Wang XJ, Xie SS, Zhang DL, Wang XP, Li BQ, Ma W & Xin H (2011) A comparison of proliferative capacity and passaging potential between neural stem and progenitor cells in adherent and neurosphere cultures. International Journal of Developmental Neuroscience 29

Suraneni P, Rubinstein B, Unruh JR, Durnin M, Hanein D & Li R (2012) The Arp2/3 complex is required for lamellipodia extension and directional fibroblast cell migration. Journal of Cell Biology 197: 239–251

Tanenbaum ME, MacUrek L, Van Der Vaart B, Galli M, Akhmanova A & Medema RH (2011) A complex of Kif18b and MCAK promotes microtubule depolymerization and is negatively regulated by aurora kinases. Current Biology 21: 1356–1365

Tardif KD, Rogers A, Cassiano J, Roth BL, Cimbora DM, McKinnon R, Peterson A, Douce TB, Robinson R, Dorweiler I, et al (2011) Characterization of the cellular and antitumor effects of MPI-0479605, a small-molecule inhibitor of the mitotic kinase Mps1. Mol Cancer Ther 10: 2267–2275

Taubenberger A V., Baum B & Matthews HK (2020) The Mechanics of Mitotic Cell Rounding. Front Cell Dev Biol 8: 1–16

Taverna E & Huttner WB (2010) Neural progenitor nuclei IN motion. Neuron 67: 906–914

Thornton GK & Woods CG (2009) Primary microcephaly: do all roads lead to Rome? Trends in Genetics 25: 501–510

Vargas-Hurtado D, Brault JB, Piolot T, Leconte L, Da Silva N, Pennetier C, Baffet A, Marthiens V & Basto R (2019) Differences in Mitotic Spindle Architecture in Mammalian Neural Stem Cells Influence Mitotic Accuracy during Brain Development. Current Biology 29: 2993–3005.e9

Viais R, Fariña-Mosquera M, Villamor-Payà M, Watanabe S, Palenzuela L, Lacasa C & Lüders J (2021) Augmin deficiency in neural stem cells causes p53-dependent apoptosis and aborts brain development. Elife 10: 1–25

Villalba A, Götz M & Borrell V (2021) The regulation of cortical neurogenesis. Curr Top Dev Biol 142: 1–66

Vink H (1971) Precision measurements of osmotic pressure in concentrated polymer solutions. Eur Polym J 7: 1411–1419

Vitre B, Gudimchuk N, Borda R, Kim Y, Heuser JE, Cleveland DW & Grishchuk EL (2014) Kinetochore-microtubule attachment throughout mitosis potentiated by the elongated stalk of the kinetochore kinesin CENP-E. Mol Biol Cell 25: 2272–2281

Walczak CE, Zong H, Jain S & Stout JR (2016) Spatial regulation of astral microtubule dynamics by Kif18B in PtK cells. Mol Biol Cell 27: 3021–3030

Weaver BAA, Bonday ZQ, Putkey FR, Kops GJPL, Silk AD & Cleveland DW (2003) Centromere-associated protein-E is essential for the mammalian mitotic checkpoint to prevent aneuploidy due to single chromosome loss. Journal of Cell Biology 162: 551–563

Yamada A, Vignes M, Bureau C, Mamane A, Venzac B, Descroix S, Viovy JL, Villard C, Peyrin JM & Malaquin L (2016) In-mold patterning and actionable axo-somatic compartmentalization for on-chip neuron culture. Lab Chip 16: 2059–2068

Zlotek-Zlotkiewicz E, Monnier S, Cappello G, Le Berre M & Piel M (2015) Optical volume and mass measurements show that mammalian cells swell during mitosis. Journal of Cell Biology 211: 765–774

