## Supplemental Figures and legends for "Brain biomechanics governs mitotic fidelity of embryonic neural progenitors"

#### Supplementary figure legends

##### Figure S1: Analysis of aRG cell density as neurogenesis progresses

(A) Representative images of *en face* views of whole-mount cerebral cortex explant at E13.5 (top) and E16.5 (bottom) showing plasma membrane stained with N-cadherin antibodies (grey). (B-C) Box and whiskers plot showing the aRG apical process density (B) and area (C) as indicated. (D) Images showing the methods used to quantify mitotic aRG cell volumes. Lateral (xz, left panel) and *en face* (xy, right panels) views of a Sox2-positive aRG mitosis. Dotty red lines indicate examples of measured areas at different z positions. Data were submitted to normality and lognormality tests for the choice of appropriate statistical test. Statistical significance was assessed with an unpaired t-test (B and C).

##### Figure S2: Mitotic timing does not account for differences between 3D and 2D aRG primary cultures

(A) Schematic representation of the protocol used to establish primary Sox2-positive aRG cultures from E13.5 dorsal telencephalon. (B) 3D aRG cells under the form of a floating sphere labelled with antibodies that recognize the stem cell marker Sox2 (red). Actin (green) and DNA (blue). (C) Box and whiskers plot showing mitotic time elapsed between nuclear envelope breakdown and anaphase onset as indicated. (D) Graph bar showing the normalized proportion of cells showing lagging chromosomes as indicated. (E) Box and whiskers plot showing the mitotic timing as indicated. (F-I) Graph bars showing the frequency of cells with lagging chromosomes as indicated. Data were submitted to normality and lognormality tests for the choice of appropriate statistics. Statistical significance was assessed with Mann-Whitney tests (C and E), chi-square test on normalized contingency data in D, ratio paired t tests (F-I).

##### Figure S3: Analysis of MT depolymerization-repolymerization assays in 2D and 3D aRG cultures

(A) Top, schematic diagram of MT depolymerization-repolymerization assays. On the bottom, representative pictures (left) and corresponding LUT showing the parameters extracted for the analysis of the number of EB1 comets and the distance travelled. (B-C) Box and whiskers plots showing the cumulative number of EB1 comets (B) and distance travelled (C) per pole during

1.5min MT repolymerization as indicated. (D) Box and whiskers plot showing the aspect ratios of 2D aRG cells plated on FN or Ncad-Fc substrates. (E-F) Box and whiskers plots showing the mean number of EB1 comets (E) and the distance travelled from the pole (F) as indicated. (G) Box and whiskers plot showing the mitotic timing in the presence of DMSO or UMK57 agonist (UMK) as indicated. (H) Graph bar showing the frequency of lagging chromosomes in the same conditions as indicated. (I) Box and whiskers plot showing the mitotic timing as indicated. (J) Graph bar showing the frequency of lagging chromosomes as indicated. Data were submitted to normality and lognormality tests for the choice of appropriate statistics. Statistical significance was assessed with unpaired t test (B), Mann-Whitney tests (C, D-F, G, I), ratio paired t test (H) and paired t test (J).

**Figure S4: Analysis of 2RG primary cultures with Dextran**

(A-B) Box and whiskers plot and bar graph showing the mitotic timing (A) and frequency of lagging chromosomes (B) in the absence (FN substrate only, FN) or presence of dextran as indicated. (C-D) Box and whiskers plot and bar graph showing the mitotic timing (C) and frequency of lagging chromosomes (D) as indicated. Data were submitted to normality and lognormality tests for the choice of appropriate statistical test. Statistical significance was assessed with non-parametric unpaired Mann-Whitney test (A and C) and paired t tests (B and D).

**Figure S5: Impact of biomechanical constraint release by actin depolymerization**

(A-B) Representative images of aRG mitotic cells from E13.5 (left panels) and E16.5 (right panels) cerebral cortices labelled with antibodies that recognize phosphorylated ERM (pERM) (green and grey in the bottom), N-cadherin (red) and DNA (blue). The white rectangles point to the regions chosen for line scan analysis in (B). (C) Box and whiskers plot showing pERM over N-cadherin (Ncad) FI at E13.5 and E16.5 as indicated. (D) Box and whiskers showing cortical actin FI levels in 3D aRG cells as indicated. (E) Representative pictures of 3D aRG mitotic cells expressing membrane Tomato before (time 0, t=0, left panels) or after (100 milliseconds, t=100msec, right panels) plasma membrane laser cuts as indicated. (F) Box and whiskers plot of the corresponding velocity recoil speeds as indicated. (G) Box and whiskers plot displaying 3D aRG mitotic cell volumes as indicated. (H-I) Box and whiskers plots displaying the cumulative number of EB1 comets per pole (H) and the distance between individual EB1 comets and the pole (I) after 2min MT repolymerization in DMSO (grey boxes) or LatB (orange boxes) 3D aRG cultures. (J) Box and whiskers plot showing mitotic timing in the presence of MPS1i in 3D aRG cultures co-treated with DMSO or LatB as indicated. (K-L) Bar graphs showing the frequency of lagging chromosomes scored in 3D aRG cells as indicated. Data were submitted to normality and lognormality tests for the choice of appropriate statistics. Statistical significance was assessed with Mann-Whitney tests (C, D, F-J), paired t test (K) or with ratio paired t test (L).

**Figure S6: Analysis of 2RG primary cultures after cortical actin depolymerization**

(A) Box and whiskers plot showing cortical actin FI levels in 2D aRG cells in DMSO (grey boxes) or LatB (, light red boxes). (B-C) Box and whiskers plots displaying cumulative number of EB1 comets polymerized per pole (B) and their distance from the pole (C) during 2min MT repolymerization as indicated. (D) Box and whiskers plot showing mitotic timing in 2D aRG cells in DMSO or LatB in combination with MPS1i. (E) Graph bar displaying the frequency of 2D aRG cells with lagging chromosomes in DMSO or LatB in combination with MPS1i. (F-G) Box and whiskers plots showing mitotic timing with MPS1i in 3D (F) or 2D (G) aRG cells co-treated with DMSO (grey boxes) or CK666 (red-purple boxes in (F) and pink in (G)). (H) Bar graph showing the frequency of 2D aRG cells with lagging chromosomes as indicated. (I) Box and whiskers plot showing mitotic timing in 3D aRG cultures co-treated with MPS1i and UMK57 in combination with either DMSO (grey boxes) or LatB (green-orange boxes). Data were submitted to normality and lognormality tests for the choice of appropriate statistics. Statistical significance was assessed with Mann-Whitney tests (A, C, D, F, G and I), unpaired t test (B), or with ratio paired t test (E and H).

**Figure S7: Analysis of the impact of biomechanical constraint release by actin depolymerization in E13.5 cerebral cortex explants**

(A) Representative images of views E13.5 cerebral cortices stained for actin (green) and DNA (blue) at the level of the ventricular lining after DMSO (top panel) or LatB (bottom panel) treatments. The dashed white lines highlight disruption of the neuroepithelium apical organization. (B) Representative images of EB1 comet number and distances from prometaphase spindle poles (in grey, top insets or after a fire LUT, bottom insets) after 2min MT repolymerization in the E13.5 cerebral cortex. Phospho-histone 3 staining (red) and DNA in blue. (C-D) Box and whiskers plots showing the cumulative number of EB1 comets polymerized per pole (C) and their distance from the pole (D) during 2min of MT repolymerization as indicated. (E) Representative images of *en face* view aRG cells from E13.5 cerebral cortices in metaphase (left panels) and anaphase (right panels) as indicated. (F) Bar graph showing the proportion of cells showing lagging chromosomes. Data were submitted to normality and lognormality tests for the choice of appropriate statistical test. Statistical significance was assessed with Mann-Whitney test (C), unpaired t test (D) and Fisher's exact test (F).

**Figure S1 Marthiens et al.**

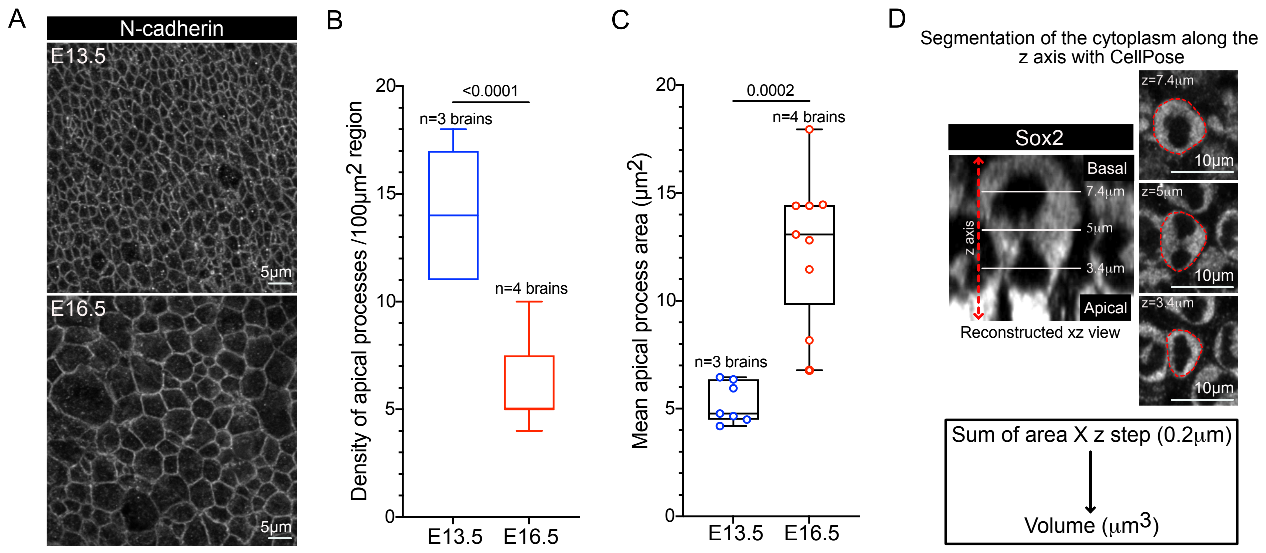

#### Figure S2 Marthiens et al.

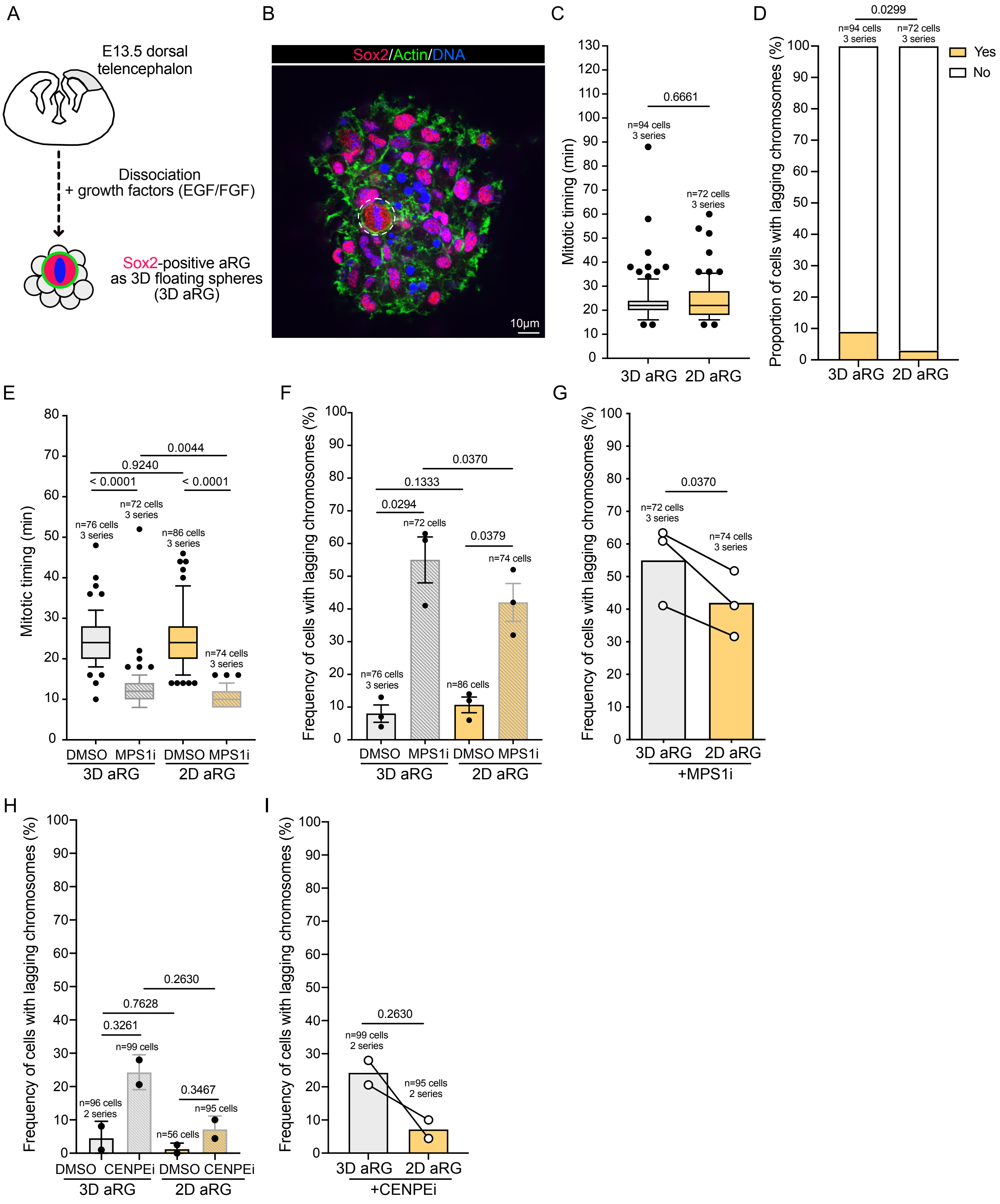

### Figure S3 Marthiens et al.

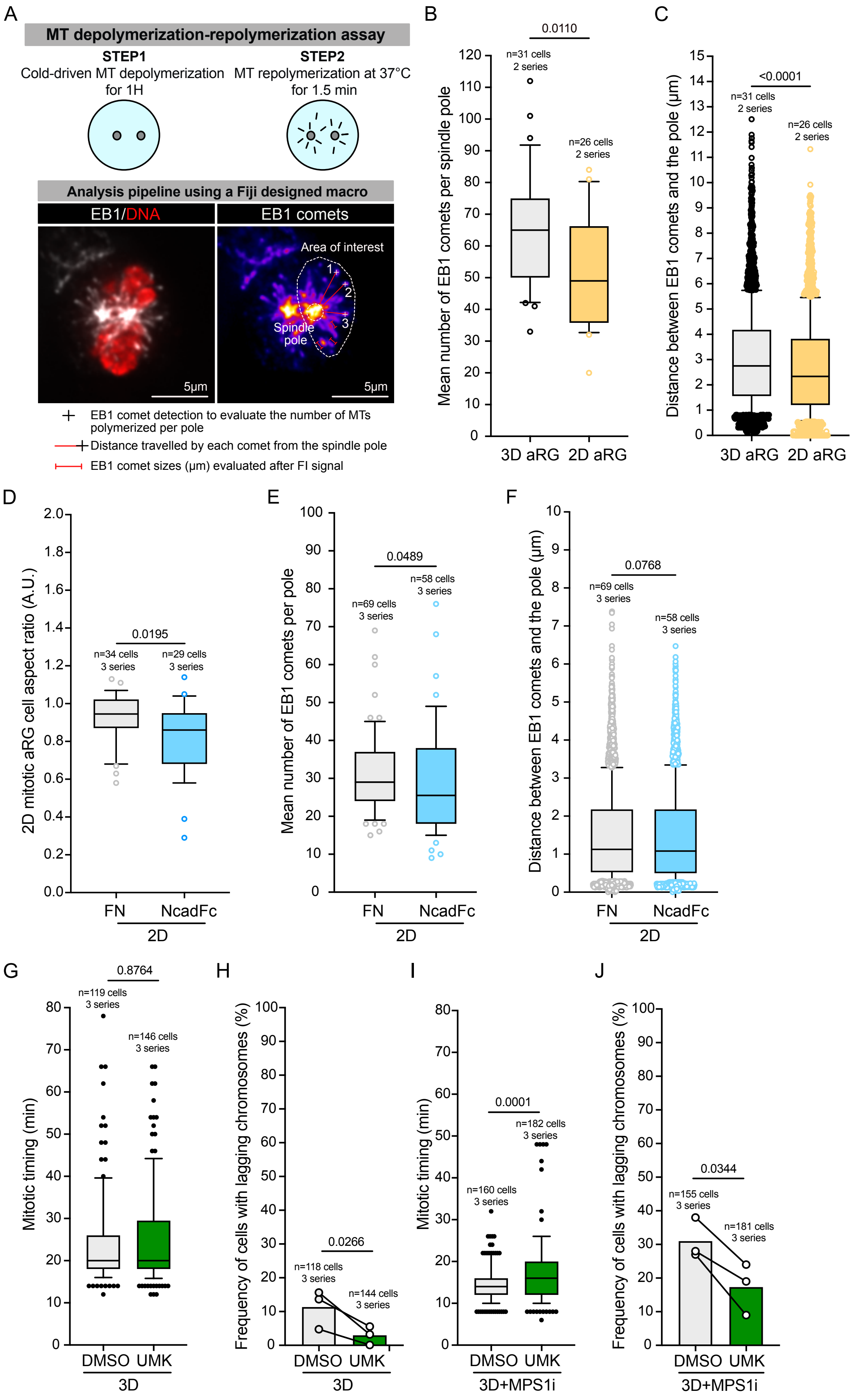

### Figure S4 Marthiens et al.

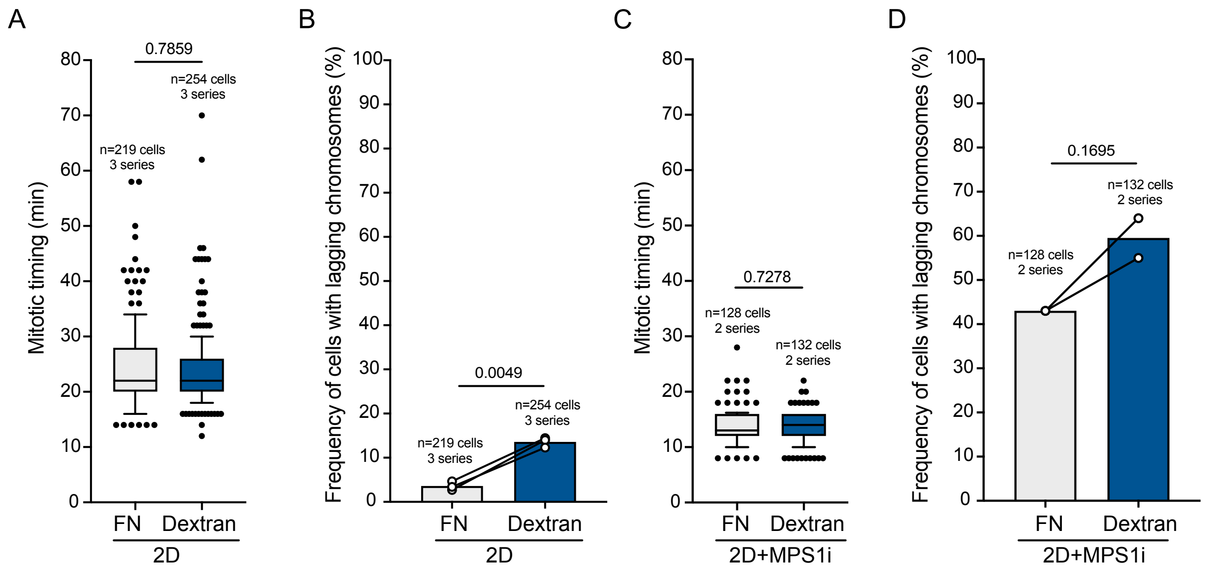

### Figure S5 Marthiens et al.

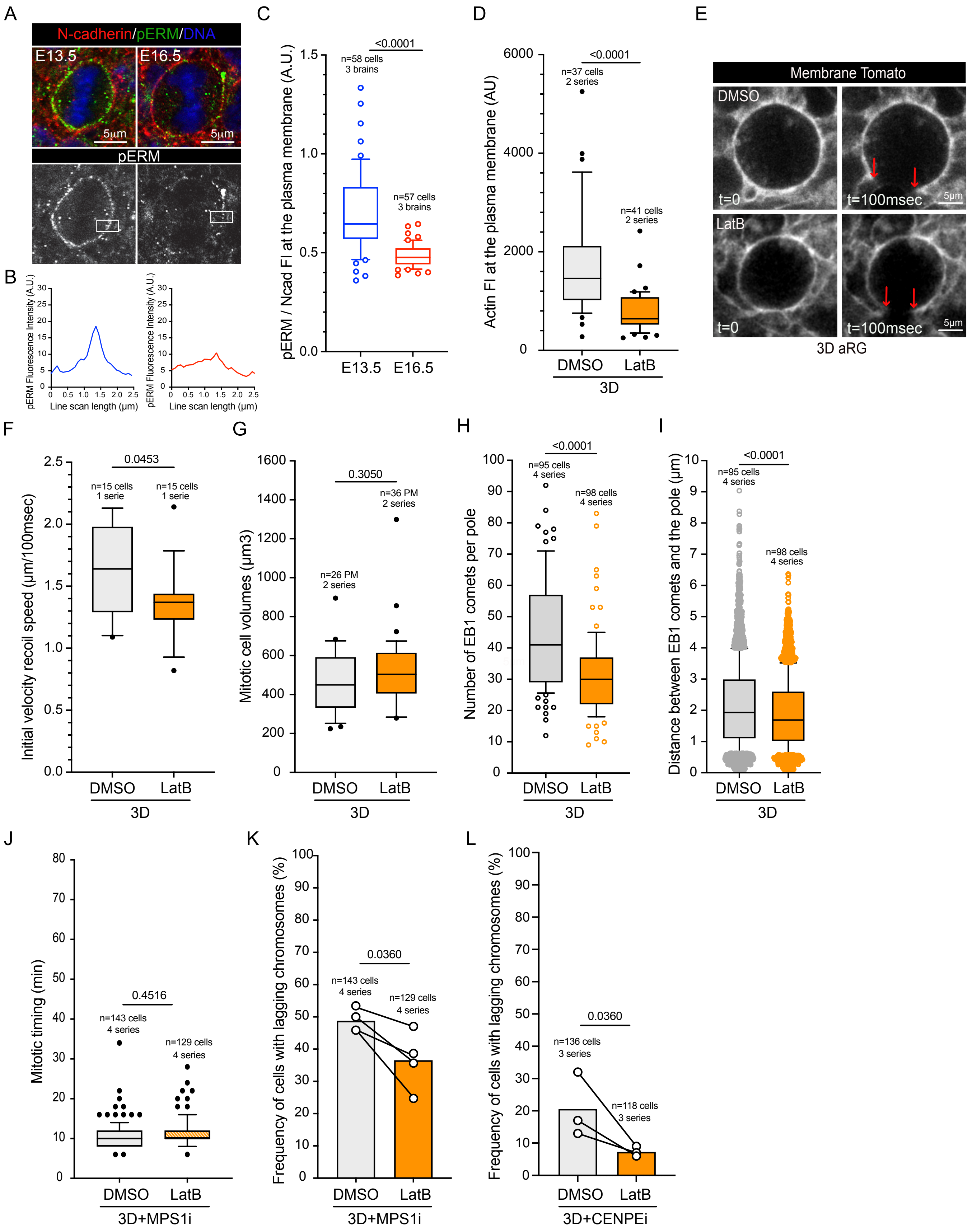

### Figure S6 Marthiens et al.

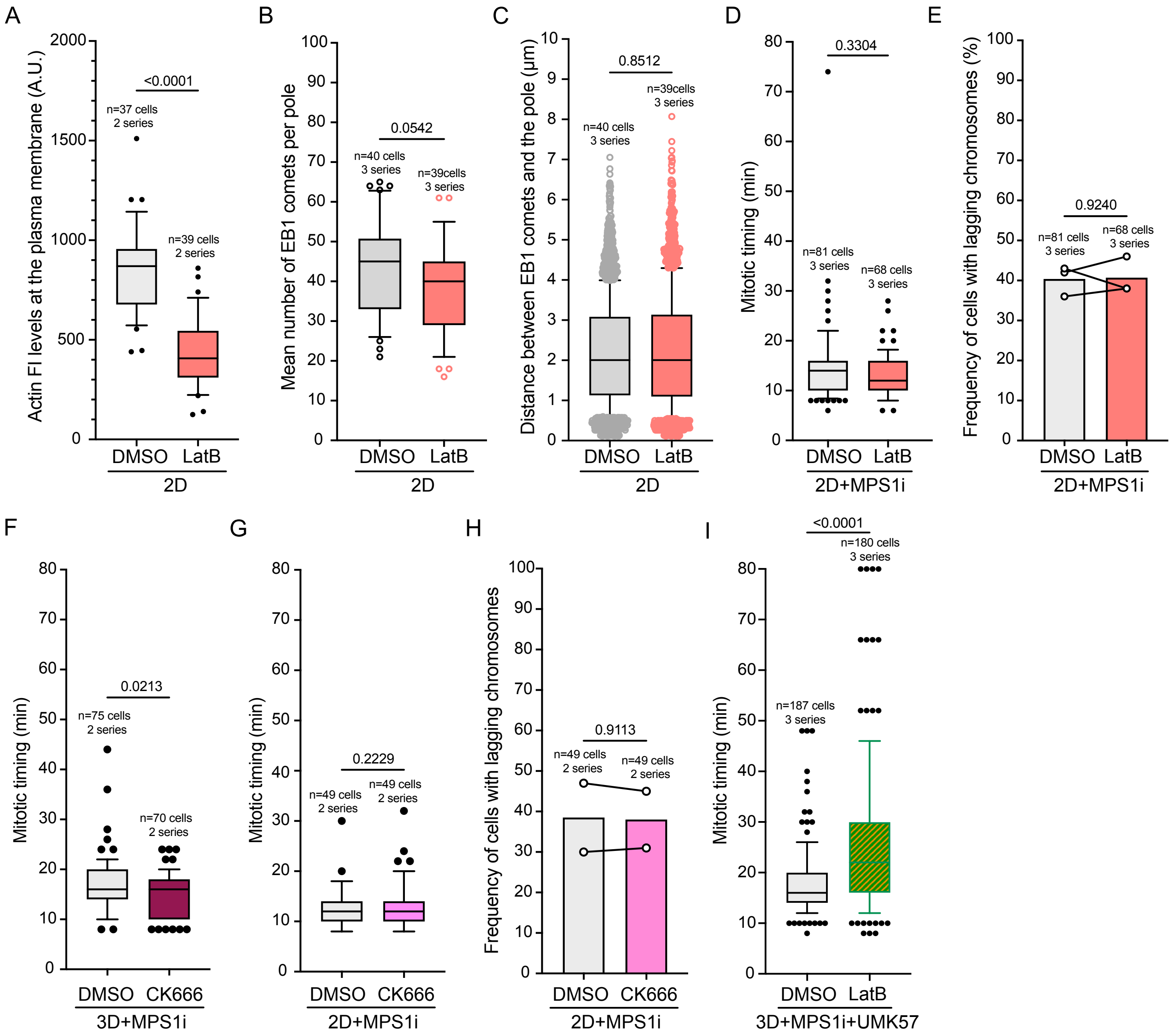

### Figure S7 Marthiens et al.

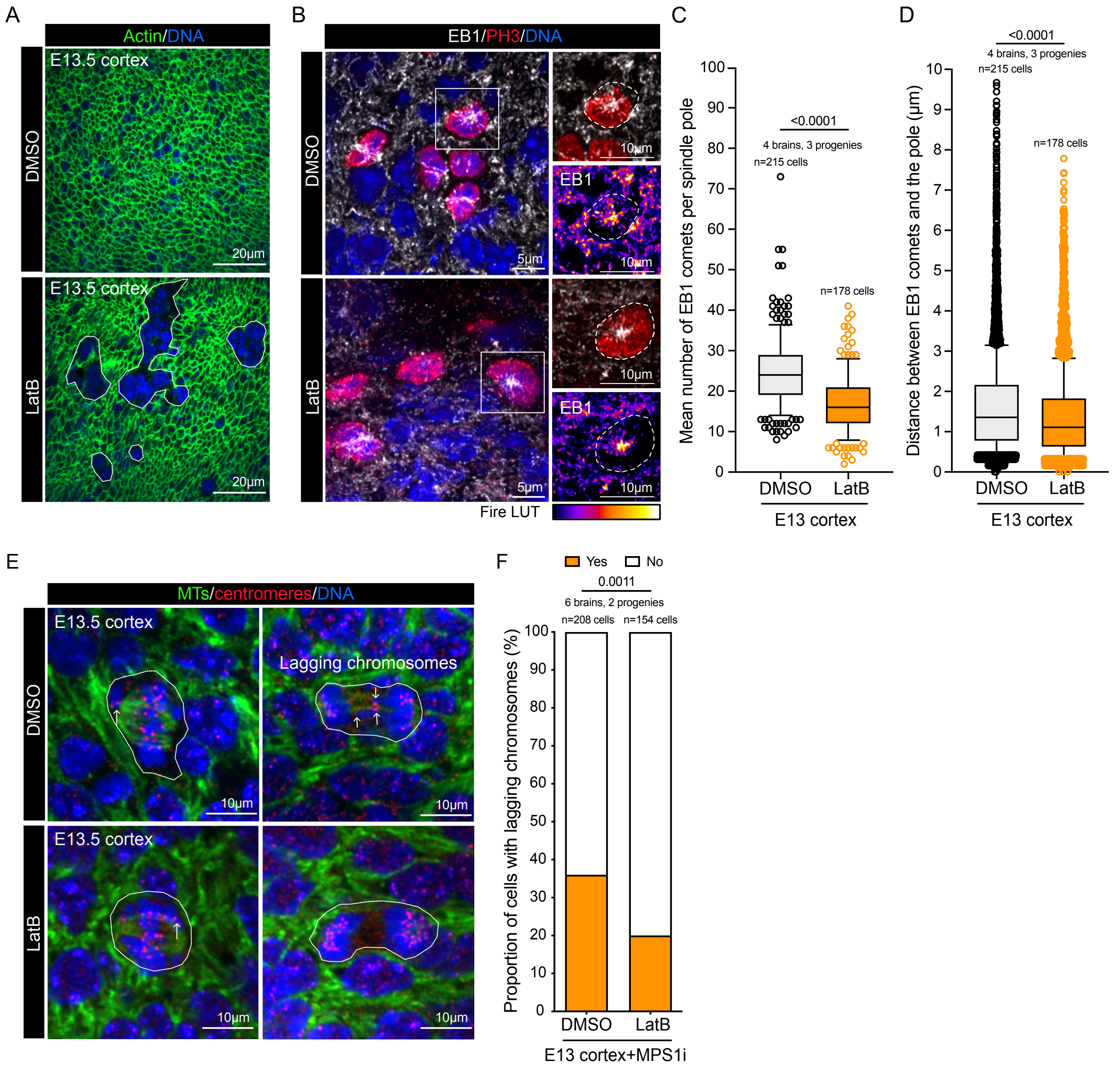
